# Distinct changes to hippocampal and medial entorhinal circuits emerge across the progression of cognitive deficits in epilepsy

**DOI:** 10.1101/2024.03.12.584697

**Authors:** Yu Feng, Keziah S. Diego, Zhe Dong, Zoé Christenson Wick, Lucia Page-Harley, Veronica Page-Harley, Julia Schnipper, Sophia I. Lamsifer, Zachary T. Pennington, Lauren M. Vetere, Paul A. Philipsberg, Ivan Soler, Albert Jurkowski, Christin J. Rosado, Nadia N. Khan, Denise J. Cai, Tristan Shuman

**Affiliations:** Icahn School of Medicine at Mount Sinai, New York, NY

## Abstract

Temporal lobe epilepsy (TLE) causes pervasive and progressive memory impairments, yet the specific circuit changes that drive these deficits remain unclear. To investigate how hippocampal-entorhinal dysfunction contributes to progressive memory deficits in epilepsy, we performed simultaneous *in vivo* electrophysiology in hippocampus (HPC) and medial entorhinal cortex (MEC) of control and epileptic mice 3 or 8 weeks after pilocarpine-induced status epilepticus (Pilo-SE). We found that HPC synchronization deficits (including reduced theta power, coherence, and altered interneuron spike timing) emerged within 3 weeks of Pilo-SE, aligning with early-onset, relatively subtle memory deficits. In contrast, abnormal synchronization within MEC and between HPC-MEC emerged later, by 8 weeks after Pilo-SE, when spatial memory impairment was more severe. Furthermore, a distinct subpopulation of MEC layer 3 excitatory neurons (active at theta troughs) was specifically impaired in epileptic mice. Together, these findings suggest that hippocampal-entorhinal circuit dysfunction accumulates and shifts as cognitive impairment progresses in TLE.

## INTRODUCTION

Temporal lobe epilepsy (TLE) is the most common form of adult-onset epilepsy^1,2^, and is often associated with cognitive impairment including deficits in spatial, verbal, and other forms of declarative memory that significantly impact quality of life^3–9^. Cognitive deficits in epilepsy are generally dissociable from chronic seizures^10,11^, suggesting that these symptoms are driven by separate neural mechanisms, yet no treatments have been developed to directly address cognitive impairment. In both people with TLE and rodent models, there is extensive cell death^12–15^ and axonal sprouting^16–19^ throughout the temporal lobe that are likely to impair normal function and lead to cognitive deficits. However, the large array of pathological changes in neuroanatomy, gene expression, and behavior emerge at different time points after an initial epileptogenic insult. Likewise, the symptoms of epilepsy^20,21^, including cognitive impairment^3–6,22^, often progressively worsen over time. These progressive changes in behavior likely reflect the continual emergence of pathological changes to the underlying neural circuits. Therefore, it is critical to understand how circuit dysfunction emerges across the timeline of epileptogenesis in order to link changes in behavior with specific circuit changes, and to develop new ways to directly treat cognitive impairment in epilepsy.

Normal cognitive function requires precisely timed neural activity in order to induce plasticity and create stable memory representations^23^. In particular, theta oscillations are strongly associated with memory processing and are thought to synchronize neural activity within and across brain regions in order to facilitate the formation of stable representations^24–35^. In rodent models of TLE, there is extensive evidence of abnormal hippocampal processing that is likely to contribute to memory impairments. For instance, in chronically epileptic rodents, CA1 place cells in hippocampus (HPC) are less precise and less stable than in Control animals^36–39^. These changes in spatial coding also coincide with an array of changes in network-wide synchronization throughout the hippocampus. In particular, epileptic rodents have decreased theta power and coherence in HPC^36,40–42^, as well as altered theta phase precession in CA1^37^ and altered theta phase locking of dentate gyrus (DG) inhibitory cells^36^. Together, these findings suggest that the synchronization of theta oscillations and neural spiking are disrupted in epileptic animals. However, it remains unclear how these network changes progress after epileptogenesis and which changes might contribute to the emergence of memory impairments.

While most studies of TLE have focused on the HPC, there are extensive anatomical changes in medial entorhinal cortex (MEC) in patients^43–47^ and in rodent models^48^. The MEC is the primary spatial input into HPC and is critical for spatial navigation and memory^27,49,50^, suggesting that it may have a prominent role in epilepsy-associated cognitive deficits. The MEC provides spatial information to HPC through grid^51^, border^52^, head direction^53^, and speed^54^ coding projections from two primary pathways: a direct input from layer 3 (MEC3) to CA1, and an indirect input from layer 2 (MEC2) stellate cells to DG^55–57^. These spatial inputs converge in the hippocampus, which creates a conjunctive representation of space and experience. This process relies on the precise timing of MEC inputs to drive plasticity^58^, and thus, a breakdown in synchronization between MEC and HPC may lead to altered spatial coding and memory performance. While one study has found an abnormal phase lag of theta oscillations between DG and MEC2^59^, no studies have examined the synchronization of theta oscillations and spike timing throughout HPC and MEC across the time course of epileptogenesis.

We previously found that there are progressive changes in the stability of CA1 spatial coding in epileptic mice after pilocarpine-induced status epilepticus (Pilo-SE), with stability deficits emerging around 6 weeks after pilocarpine treatment^36^. Notably, these changes were dissociable from the emergence of spontaneous seizures and deficits in the precision of CA1 place cells, which had already begun 3 weeks after Pilo-SE. This suggests that there are multiple pathological mechanisms driving distinct epilepsy phenotypes, with early changes producing seizures and decreased CA1 spatial precision, and later changes diminishing the stability of CA1 spatial representations. However, it remains unclear how these changes in spatial representations relate to memory performance and altered synchronization in the hippocampal-entorhinal system.

Here we used simultaneous *in vivo* electrophysiology in HPC and MEC of awake, behaving mice to determine how these key memory circuits are disrupted during the progression of memory impairments following epileptogenesis. We found distinct changes in synchronization across the hippocampal-entorhinal system that paralleled the worsening of memory deficits from 3 to 8 weeks after Pilo-SE. At the early time point, we found extensive disruptions in the synchronization of hippocampal theta and single units in epileptic mice but only minor spatial memory deficits. By the late time point, when memory impairment was more severe, we found disrupted synchronization within the MEC and between the MEC and HPC. Furthermore, within MEC, we found that a distinct population of excitatory neurons in MEC3, active near the trough of theta oscillations, were specifically disrupted in epileptic mice. Together, these results suggest that the severity of spatial memory impairments is paralleled by the severity of network-wide communication within the MEC and between the MEC and HPC.

## RESULTS

### Spatial memory deficits in epileptic mice progressively worsen from 3 to 8 weeks after Pilo-SE, and this progression cannot be explained by seizures or cell loss in HPC or MEC

Spatial memory deficits are well-established in mouse models of TLE^40,60–63^, but are typically examined many weeks after epileptogenesis, preventing examination of the progression of behavioral changes. Therefore, we first set out to establish the time course of spatial memory deficits in the pilocarpine mouse model of chronic TLE. We previously found that epileptic mice have impaired spatial coding in CA1, with early deficits in spatial precision (i.e., information content) already at 3 weeks after Pilo-SE and late-onset deficits in spatial stability^36^ emerging around 6-8 weeks after Pilo-SE. We therefore performed a novel object location (NOL) task in Control and Epileptic mice either 3 or 8 weeks after Pilo-SE (Figure 1A). The NOL task is a well-established measure of spatial memory^64,65^ that is dependent on both HPC^66^ and MEC^67^.

**Figure 1:**
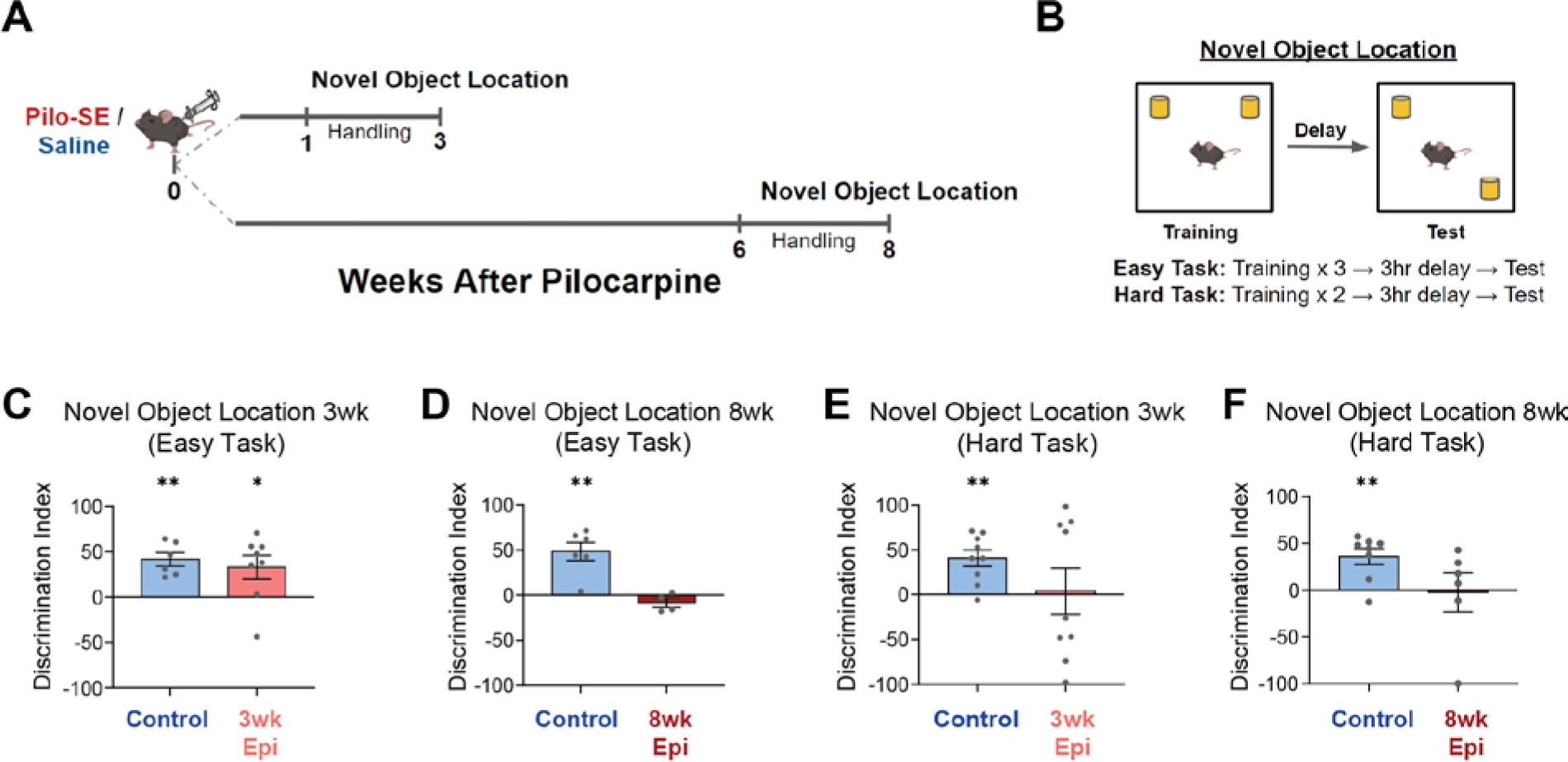
Early-onset and progressive spatial memory deficits after pilocarpine-induced status epilepticus (Pilo-SE) **A.** Timeline of behavioral testing for the novel object location (NOL) task. **B.** NOL training was performed using an Easy (three 6-min training trials) or Hard (two 6-min training trials) version of the task. Testing occurred 3 hours after the final training sessions. **C.** On the Easy version of the NOL task both Control and 3wk Epileptic (3wk Epi) groups demonstrated a significant preference for the moved object (one-sample t-test: Control: *p* < 0.01, N = 6; 3wk Epileptic: *p* < 0.05, N = 8). **D.** At 8 weeks after Pilo-SE, the Control mice showed a preference for the moved object on the easy NOL task, while the 8wk Epileptic (8wk Epi) mice showed no preference (one-sample t-test: Control: *p* < 0.01, N = 6; 3wk Epileptic: *p* > 0.05, N = 4). **E-F**. On the Hard NOL task, neither the 3wk (E) nor 8wk (F) Epileptic mice showed a preference to investigate the moved object, while age-matched Control mice showed a significant preference to investigate the moved object (**E**, one-sample t-test: Control: *p* < 0.01, N = 9; 3wk Epileptic: *p* > 0.05, N = 9; **F**, one-sample t-test: Control: *p* < 0.01, N = 8; 3wk Epileptic: *p* > 0.05, N = 6). Error bars represent s.e.m. *p<0.05, **p<0.01.

Animals were first exposed to two identical objects in a 1ft x 1ft arena. After a 3-hour delay, they were placed back in the same arena with one object moved to a new location. We used two separate training protocols in order to vary the difficulty of the task. In the easier version of this task (labeled Easy), the training session consisted of three 6-min back-to-back exposures (between each exposure animals were removed, the chamber was cleaned, and the animals were immediately returned to the arena), while the more challenging version (labeled Hard) consisted of only two 6-min exposures. Spatial memory was assessed by the discrimination index (DI), which represents the preference to investigate the moved object over the unmoved object (Figure 1B, see Methods and Materials).

Using the Easy NOL task, we found progressive memory deficits that emerged between 3 and 8 weeks after Pilo-SE. At 3 weeks after Pilo-SE, both Control and Epileptic animals showed a preference for the moved object (as a measure of memory for the prior object location) (Figure 1C). However, by 8 weeks after Pilo-SE, the Epileptic animals no longer showed a preference for the moved object (Figure 1D). Thus, on this Easy NOL task, Epileptic mice showed progressively worse performance from 3 to 8 weeks after Pilo-SE.

We next increased the difficulty of the NOL task to probe for more subtle changes in memory (Figure 1B). In this Hard NOL task, Control mice showed a preference for the moved object, but Epileptic mice at both 3 and 8 weeks after Pilo-SE did not show a preference (Figure 1E-F).

Raw exploration times of both objects during training were not significantly different between Epileptic and Control mice, and exploration times during test sessions were consistent with the DI metric (Figure S1). Together, these results indicate that spatial memory deficits in Epileptic mice are relatively subtle at the 3-week time point and progressively worsen between 3 and 8 weeks after Pilo-SE.

Neuronal cell death is well-established in chronic epilepsy^68,69^, and it is possible that the progressive changes in memory performance could reflect ongoing cell death. To better characterize the time course of neurodegeneration and cell loss, we performed FluoroJade C (FJC, a marker of degenerating neurons) staining^70^ and NeuN immunohistochemistry (a neuronal marker)^71^ in the HPC and MEC at 2 days, 3 weeks, or 8 weeks after Pilo-SE. We found extensive FJC expression 2 days after Pilo-SE in all areas examined, including CA1, DG, MEC2, and MEC3 (Figure S2A-H), suggesting widespread neural degenerative signaling in the days after Pilo-SE. However, we found minimal FJC staining at 3 weeks or 8 weeks after Pilo-SE in Epileptic mice (Figure S2A-H), indicating that neurodegeneration primarily occurs within 2 days after Pilo-SE. This was further confirmed by NeuN immunohistochemistry, which showed significantly reduced NeuN expression in the dentate hilus and the ventral portion of MEC3 beginning 2 days after Pilo-SE that remained at a similar level across the 3-week and 8-week time points (Figure S2I-M). We did not find significantly reduced NeuN staining in the dorsal portion of MEC2 or MEC3 in Epileptic mice, suggesting that the high levels of FJC staining (Figure S2A-H) do not necessarily lead to permanent cell loss in these areas (Figure S2I-M). This disconnect between FJC and NeuN expression indicates that FJC staining is driven by early degenerative processes that could recover within the first weeks after Pilo-SE. Together, these data indicate that most neuronal cell loss in the HPC and MEC of Epileptic mice occurs within 2 days of Pilo-SE and is unlikely to directly drive the progression of memory deficits between 3 and 8 weeks after Pilo-SE (Figure 1C,D).

We also found that progressive memory deficits (Figure 1C,D) are dissociable from seizure frequency as we observed no differences in seizure frequency between Epileptic mice at 3 or 8 weeks after Pilo-SE (see Figure 2B). This suggests that progressive memory impairments have distinct neural correlates from the onset and frequency of seizure events. Thus, we next sought to determine how activity within and between HPC and MEC circuits changes from 3 to 8 weeks after Pilo-SE in Control and Epileptic mice.

**Figure 2:**
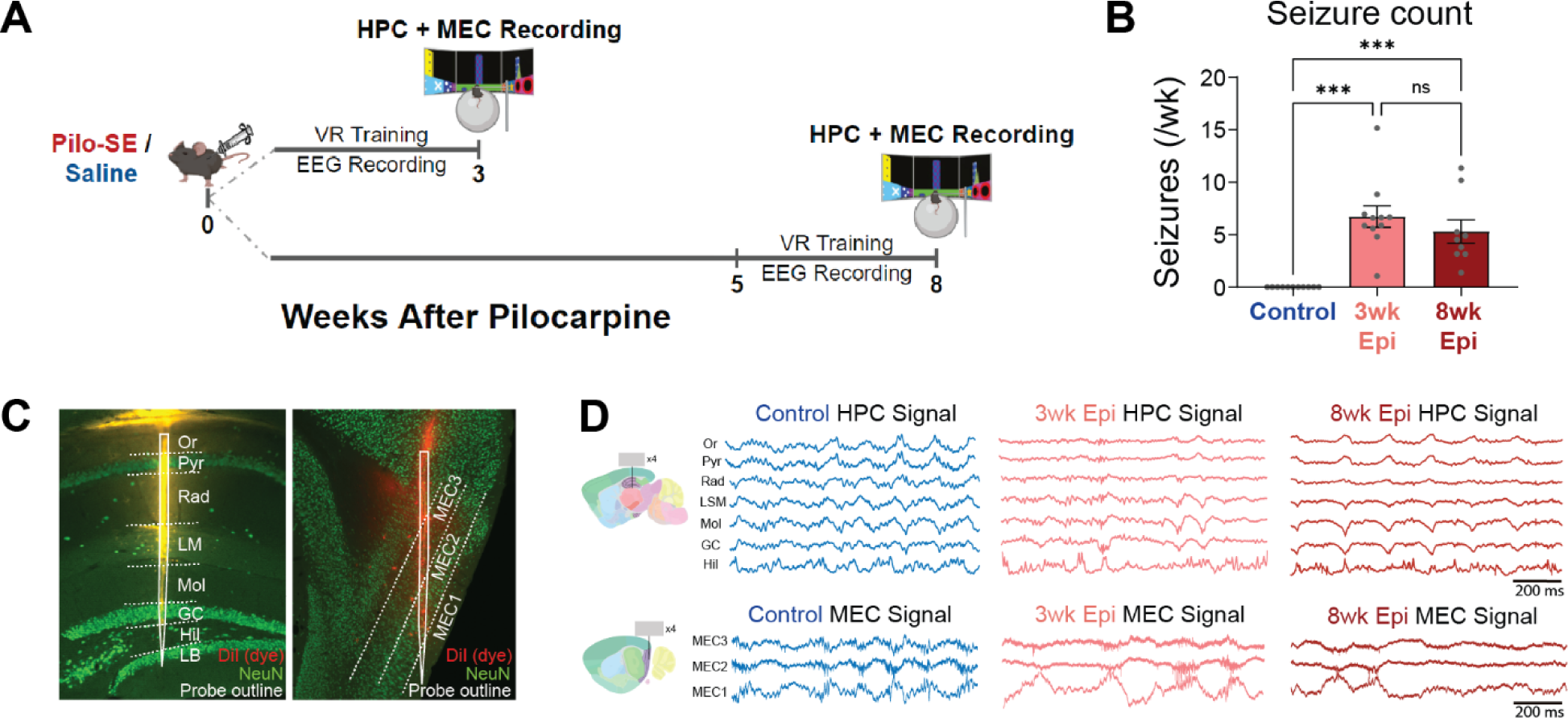
Simultaneous *in vivo* electrophysiology of HPC and MEC in Control and Epileptic mice 3 or 8 weeks after Pilo-SE. **A.** Schematic of experimental timeline. Simultaneous hippocampus (HPC) and medial entorhinal cortex (MEC) recordings were performed in head-fixed mice 3 or 8 weeks after Pilo-SE or saline injection. During recordings, animals ran through a virtual linear track for water rewards. **B.** Chronic wireless EEG prior to silicon probe recordings showed seizure frequency was not significantly different between 1-3 and 6-8 weeks after Pilo-SE (N = 11, Control; N = 11, 3wk Epileptic; N = 9, 8wk Epileptic; one-way ANOVA, *F* = 18.9, *p* < 0.0001; post hoc tests corrected by Bonferroni, Control v. 3wk Epileptic: *p* < 0.0001; Control v. 8wk Epileptic: *p* < 0.001; 3wk Epileptic v. 8wk Epileptic: *p* = 0.74). **C.** Silicon probes were inserted into dorsal HPC (spanning CA1 and dentate gyrus) and MEC (spanning from MEC3 to MEC1). Probes were covered with DiI prior to recordings to facilitate later visualization of the probe tracts. **D.** Example signals from each layer of HPC and MEC during recording in Control, 3wk Epileptic, and 8wk Epileptic groups. Or, stratum oriens; Pyr, stratum pyramidale; Rad, stratum radiatum; LM, stratum lacunosum moleculare; Mol, molecular layer of dentate gyrus; GC, granule cell layer; Hil, hilus of dentate gyrus; LB, lower blade of granule cell layer. Error bars represent s.e.m. *p<0.05, ***p<0.001.

### Early onset desynchronization in HPC of Epileptic mice

To determine how synchronization of the hippocampal-entorhinal circuit is disrupted across the progression of memory deficits in epilepsy, we performed *in vivo* acute extracellular electrophysiological recordings using high-density silicon probes in awake, behaving, head-fixed mice. We recorded simultaneously from the HPC and MEC of Control and Epileptic mice at 3 or 8 weeks after Pilo-SE as mice navigated a virtual linear track (Figure 2A). We also performed chronic wireless electroencephalogram (EEG) recordings for three weeks prior to silicon probe recordings to examine the onset of chronic seizures. All pilocarpine-treated mice showed spontaneous seizures throughout the EEG recording period and we found no differences in the frequency of seizures recorded in the 3wk Epileptic or 8wk Epileptic mice (Figure 2B). No seizures were observed in Control mice. To engage spatial processing during silicon probe recordings, mice were head-fixed atop a Styrofoam ball and trained to run through a virtual reality (VR) linear track for water rewards^36^. After animals were well-trained (8-12 total sessions), we performed dual-region acute electrophysiology recordings with two 256-channel silicon probes (4 shanks with 64 channels per shank) by lowering one probe into the dorsal HPC and the other into the superficial layers of MEC (Figure 2C). Example local field potential (LFP) traces for each subregion in HPC and MEC are shown in Figure 2D and probe tracts from each animal are shown in Figure S3. To isolate hippocampal processing, we limited our analysis to periods of locomotion. We found no differences in running speed between any of the groups (Figure S4A), consistent with our previous work^36^. In addition, Control mice in the 3wk and 8wk groups had no differences in theta power (Figure S4B), or theta coherence (Figure S4C,D) and were therefore combined for all further analyses.

We first examined the time course of intra-HPC synchronization deficits, which have been previously found in multiple models of epilepsy^36,42^. We measured theta power and coherence along the CA1-DG axis, as well as the theta phase locking (the propensity of a neuron to be active at a specific phase of theta oscillations^36^) of inhibitory neurons within HPC. Together, these measures reflect the degree of coordination between long-range inputs and local neural activity in HPC^72,73^. At both 3 weeks and 8 weeks after Pilo-SE, Epileptic animals showed reduced theta power in the lacunosum moleculare (LM) layer of the CA1 region and the molecular (Mol) layer of the DG (Figure 3A). These sublayers correspond to the inputs from entorhinal cortex and reduced power suggests reduced input strength or coordination in Epileptic mice. We next examined HPC theta coherence across each recorded channel pair. At 3 weeks after Pilo-SE we found decreased theta coherence between CA1 and DG as well as within the DG (Figure 3B), with the most prominent reductions occurring between the DG hilus (Hil) and the CA1 stratum oriens (Or), CA1 lacunosum moleculare (LM), DG molecular layer (Mol), and DG granular cell layer (GC). By 8 weeks after Pilo-SE, we found additional deficits in coherence between the DG hilus and CA1 pyramidal layer (Pyr) (Figure 3B). These theta coherence deficits reflect reduced coordination of hippocampal processing across CA1 and DG in Epileptic mice. Importantly, both deficits in hippocampal theta power and coherence primarily emerged early after epileptogenesis, within 3 weeks of Pilo-SE, which aligned with early-onset relatively minor memory impairment on the NOL task (Figure 1).

**Figure 3:**
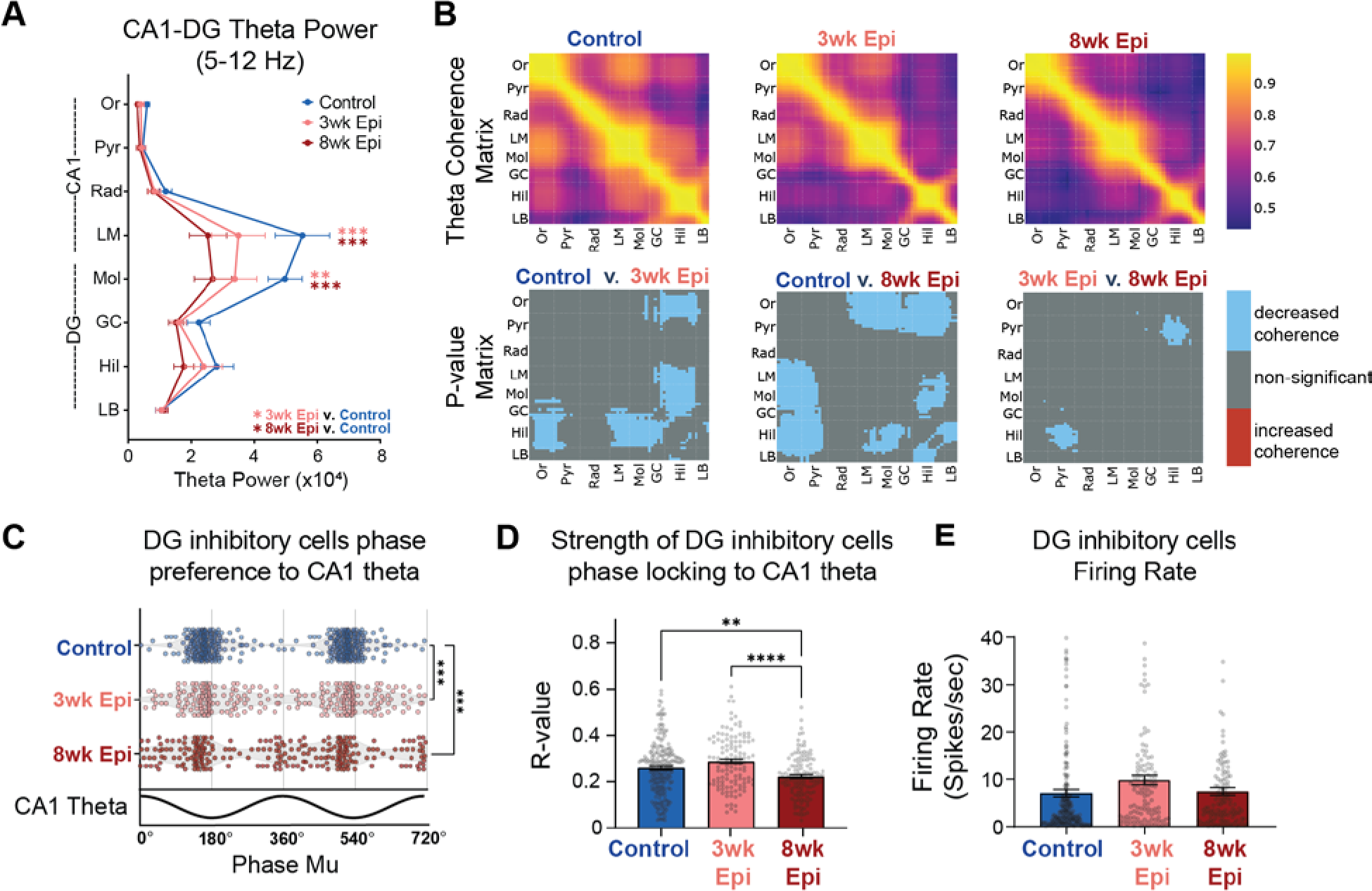
Deficits in HPC synchronization primarily emerge by 3 weeks after Pilo-SE. **A.** Theta power recorded from each hippocampus layer. Epileptic animals had reduced power in CA1 lacunosum moleculare (LM) and dentate gyrus (DG) molecular layer (Mol) in both 3wk and 8wk Epileptic groups (N = 11, Control animals; N = 11, 3wk Epileptic animals; N = 11, 8wk Epileptic animals; Repeated-measures mixed-effects model, *F*Group x Region (14,204) = 3.1 *p* < 0.001; Holm-Sidak corrected: LM: Control v. 3wk Epileptic *p* < 0.05; Control v. 8wk Epileptic *p* < 0.001; Mol: Control v. 3wk Epileptic *p* < 0.01; Control v. 8wk Epileptic *p* < 0.0001). **B.** Theta coherence between each channel pair along the probe in HPC in Control (top left), 3wk Epileptic (top middle) and 8wk Epileptic (top right) mice. P value matrix (bottom row) of significant changes in coherence at each location between experimental groups (N = 11, Control animals; N = 11, 3wk Epileptic animals; N = 11, 8wk Epileptic animals; welch t-test with post-hoc correction for group comparisons by using alpha = 0.017; blue: decrease coherence; red: increase coherence). Both 3wk and 8wk Epileptic groups showed reduced theta coherence between CA1 and DG, with progressively decreased theta coherence between CA1 pyramidal (Pyr) and DG hilus (Hil) from 3wks to 8wks after Pilo-SE. **C.** Phase preference to CA1 theta for DG interneurons in Control and Epileptic animals. Each dot represents one interneuron, and data is double plotted for visualization purposes. Both 3wk Epileptic and 8wk Epileptic groups showed disrupted phase preferences compared to Control (n = 213, Control cells; n = 119, 3wk Epileptic cells; n = 123, 8wk Epileptic cells; circular k-test for equal concentrations: Control v. 3wk Epileptic *p* < 0.00003, Control v. 8wk Epileptic *p* < 0.00003; post-hoc correction for group comparisons by using alpha = 0.017). **D.** Phase locking strength (R value) of DG interneurons to CA1 theta. 8wk Epileptic group showed reduced phase locking strength compared to both Control and 3wk Epileptic group (n = 213, Control cells; n = 120, 3wk Epileptic cells; n = 125, 8wk Epileptic cells; one-way ANOVA, *F* (2, 455) = 11.5, *p* < 0.0001; Bonferroni corrected: Control v. 8wk Epileptic *p* < 0.01; 3wk Epileptic v. 8wk Epileptic *p* < 0.0001). **E.** No difference in firing rate of DG inhibitory cells (n = 213, Control cells; n = 120, 3wk Epileptic cells; n = 125, 8wk Epileptic cells; one-way ANOVA, *F* (2, 455) = 2.6, *p* = 0.07). Or, stratum oriens; Pyr, stratum pyramidale; Rad, stratum radiatum; LM, stratum lacunosum moleculare; Mol, molecular layer of dentate gyrus; GC, granule cell layer; Hil, hilus of dentate gyrus; LB, lower blade of granule cell layer; DG, dentate gyrus. Error bars represent s.e.m. *p<0.05, **p<0.01, ***p<0.001, ****p<0.0001 for A, D, E; *p<0.017, **p<0.003, ***p<0.0003 for C

One possibility is that the reduced theta power and coherence seen in Epileptic mice could have been a result of recent seizures that disrupted neural activity. To address this possibility, we examined the correlation between theta power or coherence and seizure frequency or seizure recency (the amount of time between the last seizure and the recording). We found that decreased HPC theta power and coherence in Epileptic mice was not significantly correlated with seizure frequency or recency (Figure S5A-D), suggesting that hippocampal theta power and coherence deficits are unrelated to seizure susceptibility.

We previously found that interneurons in the DG have abnormal phase locking relative to theta oscillations in Epileptic mice 17+ weeks after Pilo-SE^36^. This abnormal phase locking is independent of deficits in theta power^36^ and may contribute to abnormal spatial processing in Epileptic mice^74,75^. To examine the time course of these phase locking deficits, we first isolated single units from our recordings and identified putative interneurons based on firing properties^34,36,76^ (Figure S6, see Methods). We then calculated the mean phase of firing (mu) and phase locking strength (r) for each interneuron^36^. In Control mice, DG interneurons were reliably phase locked near the trough of CA1 theta. However, in Epileptic mice at both 3 and 8 weeks after Pilo-SE, we found an altered distribution of preferred firing phases (mu-values), with a significant reduction in the concentration of trough-preferring cells (Figure 3C). The strength of phase locking (r) was also slightly reduced in the Epileptic mice, but only at 8 weeks after Pilo-SE (Figure 3D). Notably, the firing rates of DG inhibitory neurons were unaltered either 3 or 8 weeks after Pilo-SE (Figure 3E), suggesting that the frequency of inhibitory firing was unchanged, but the timing relative to theta was disrupted. Overall, DG interneurons had disrupted theta phase locking early after epileptogenesis, primarily occurring within the first 3 weeks after Pilo-SE, similar to what we observed with HPC theta power and coherence (Figure 3A-B). Together, these data demonstrate that HPC theta synchronization and spike timing is largely disrupted within the first 3 weeks of epileptogenesis, matching the timeline of early minor memory deficits (Figure 1C,E) and the onset of seizures (Figure 2B).

### Late onset deficits in the timing of MEC excitatory inputs to HPC in Epileptic mice

Given the substantial theta synchronization deficits we observed in HPC of Epileptic mice, we next examined the timing of upstream inputs from MEC. Excitatory neurons in MEC2 and MEC3 project into HPC^55^, and are a powerful driver of HPC theta^76^. Therefore, it is possible that abnormal spike timing or reduced activity in these populations could underlie the HPC theta desynchronization we found at 3 and 8 weeks after Pilo-SE. To test this hypothesis, we isolated excitatory neurons in MEC2 and MEC3 (Figure S6) and examined their theta phase locking. Since MEC2 neurons project directly to DG, we examined the phase locking of these neurons to DG theta oscillations. At 3 weeks after Pilo-SE, we found that theta phase locking of MEC2 excitatory cells was fully intact in Epileptic animals, with no changes in the distribution of theta phase preference (Figure 4A), phase locking strength (Figure 4B), or firing rate (Figure 4C). At 8 weeks after Pilo-SE we saw similar results, with the exception of a relatively small change in the concentration of theta phase preferences (Figure 4A). Together, these results suggest that changes in MEC2 theta phase locking are unlikely to be contributing to the early theta desynchronization observed in HPC at 3 weeks after Pilo-SE.

**Figure 4:**
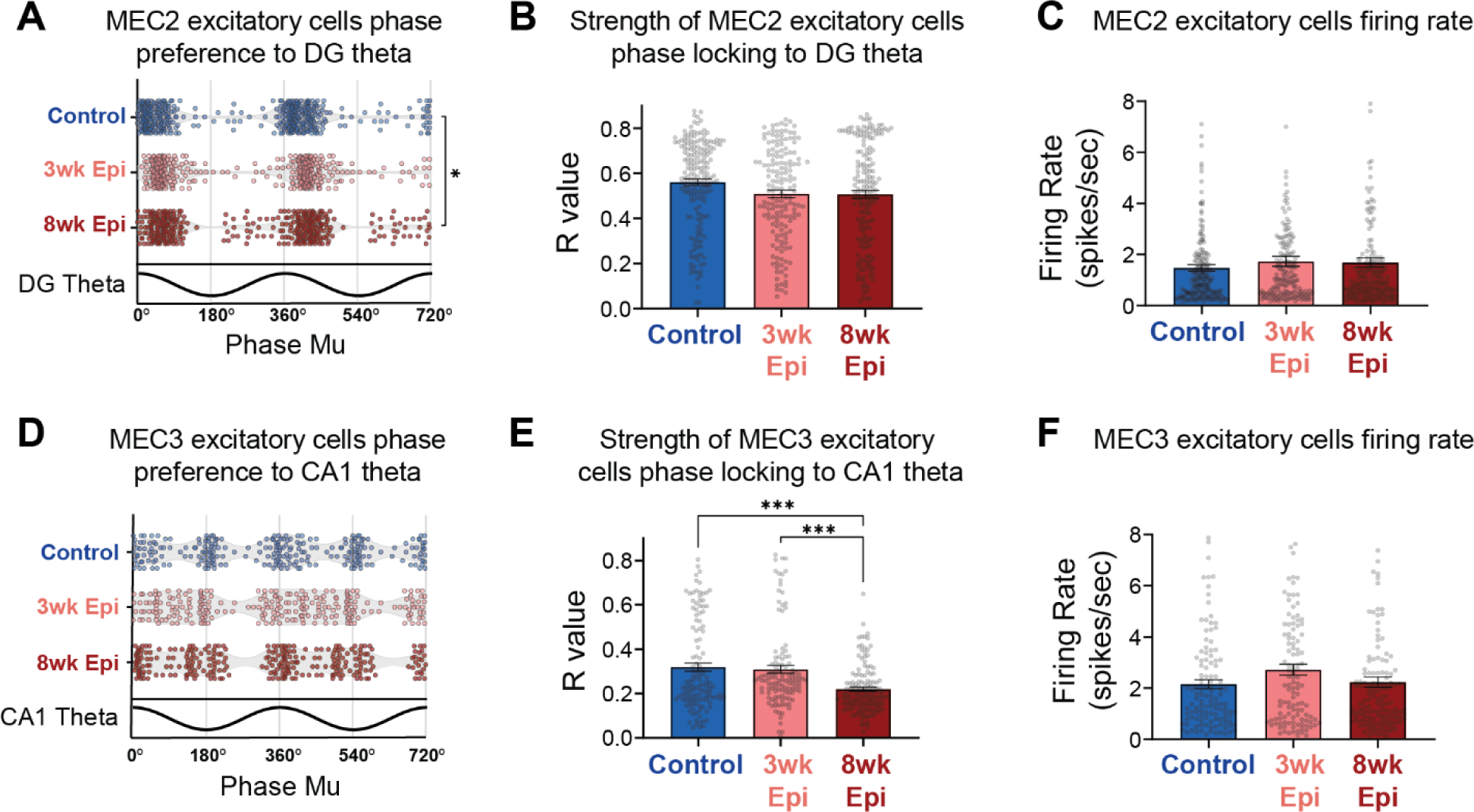
MEC3 single units show late-onset reduction in theta phase locking strength. **A.** Phase preference to DG theta for MEC2 excitatory cells in Control and Epileptic animals (n = 211, Control cells; n = 173, 3wk Epileptic cells; n = 186, 8wk Epileptic cells; circular k-test for equal concentrations: Control v. 8wk Epileptic *p* < 0.017; post-hoc correction for group comparisons by using alpha = 0.017). **B.** No changes in phase locking strength of MEC2 excitatory cells to DG theta in Epileptic mice (n = 214, Control cells; n = 175, 3wk Epileptic cells; n = 188, 8wk Epileptic cells; one-way ANOVA, *F* (2, 574) = 1.8, *p* > 0.05). **C.** No changes in firing rate of MEC2 excitatory cells in Epileptic mice (n = 214, Control cells; n = 175, 3wk Epileptic cells; n = 188, 8wk Epileptic cells; one-way ANOVA, *F* (2, 574) = 0.6, *p* > 0.05). **D.** No changes in phase preference to CA1 theta for MEC3 excitatory cells in Control and Epileptic animals (n = 121, Control cells; n = 120, 3wk Epileptic cells; n = 135, 8wk Epileptic cells; circular k-test for equal concentrations: *p* > 0.017 for all comparisons; post-hoc correction for group comparisons by using alpha = 0.017). **E.** Reduced phase locking strength of MEC3 excitatory cells to CA1 theta in Epileptic mice 8 weeks after Pilo-SE (n = 122, Control cells; n = 122, 3wk Epileptic cells; n = 137, 8wk Epileptic cells; one-way ANOVA, *F* (2, 378) = 14.6, *p* < 0.0001; Bonferroni corrected: Control v. 8wk Epileptic *p* < 0.0001, 3wk Epileptic v. 8wk Epileptic *p* < 0.0001). **F.** No changes in firing rate of MEC3 excitatory cells in Epileptic mice (n = 122, Control cells; n = 122, 3wk Epileptic cells; n = 137, 8wk Epileptic cells; one-way ANOVA, *F* (2, 378) = 1.8, *p* > 0.05). Error bars represent s.e.m. *p<0.05, **p<0.01, ***p<0.001 for B,C,E,F; *p<0.017 for A, D

To test whether changes in MEC3 excitatory neurons, which project directly to CA1, were contributing to the theta desynchronization seen in HPC, we next examined the phase locking of MEC3 excitatory neurons to CA1 theta. We found no significant changes in the distribution of phase preferences at either 3 or 8 weeks after Pilo-SE (Figure 4D). However, we did find a late-onset reduction in phase locking strength in Epileptic mice at 8 weeks after Pilo-SE (Figure 4E), suggesting a breakdown in the precise timing of MEC3 excitatory neuron spiking during the progression of epileptogenesis. No firing rate changes were detected in this population at either time point (Figure 4F), again suggesting that overall activity levels were maintained, but that changes were occurring primarily in the timing of this neural activity. Together, these results indicate that MEC excitatory activity in both layers 2 and 3 was intact at the early time point (3 weeks after Pilo-SE) when HPC deficits and minor memory impairment were already observed and are therefore unlikely to drive these deficits. However, we did observe late-onset MEC phase locking deficits, primarily in the strength of phase locking in MEC3 excitatory neurons to downstream CA1 theta, that match the time course of progressive memory deficits. Notably, these deficits also align with the time course of reduced stability of place cells in CA1 of Epileptic mice^36^. Given the extensive evidence that the timing of direct entorhinal inputs to CA1 can mediate plasticity of CA1 neurons^77–80^, this suggests that disrupted timing of MEC3 excitatory neurons may be an important mediator of CA1 instability and memory deficits in epilepsy.

### Deficits in MEC3 spike timing are driven by a distinct subpopulation that is preferentially active at the trough of MEC theta

While MEC3 excitatory neurons are often thought to be a homogenous population, there is some evidence of distinct subpopulations that are active at different phases of theta oscillations^33^. Indeed, we observed two clusters of phase preferences in MEC3 excitatory neurons in all groups of animals (Figure 4D). To gain more insights into these subpopulations of MEC3 excitatory cells, we performed k-means clustering on their preferred phase (mu) and phase locking strength (r) to the local MEC theta and these cells fell into two distinct clusters (Figure 5A). Within each experimental group, the clusters reliably formed two populations, with one active primarily at the theta trough (Trough-locked) and one primarily active at the thet apeak (Peak-locked). We found no difference in the spatial distribution of these clusters along the silicon probe (Figure 5A), indicating that these two cell populations are evenly dispersed throughout MEC3 and that this effect is not due to cells being picked up from neighboring regions above or below MEC3.

**Figure 5:**
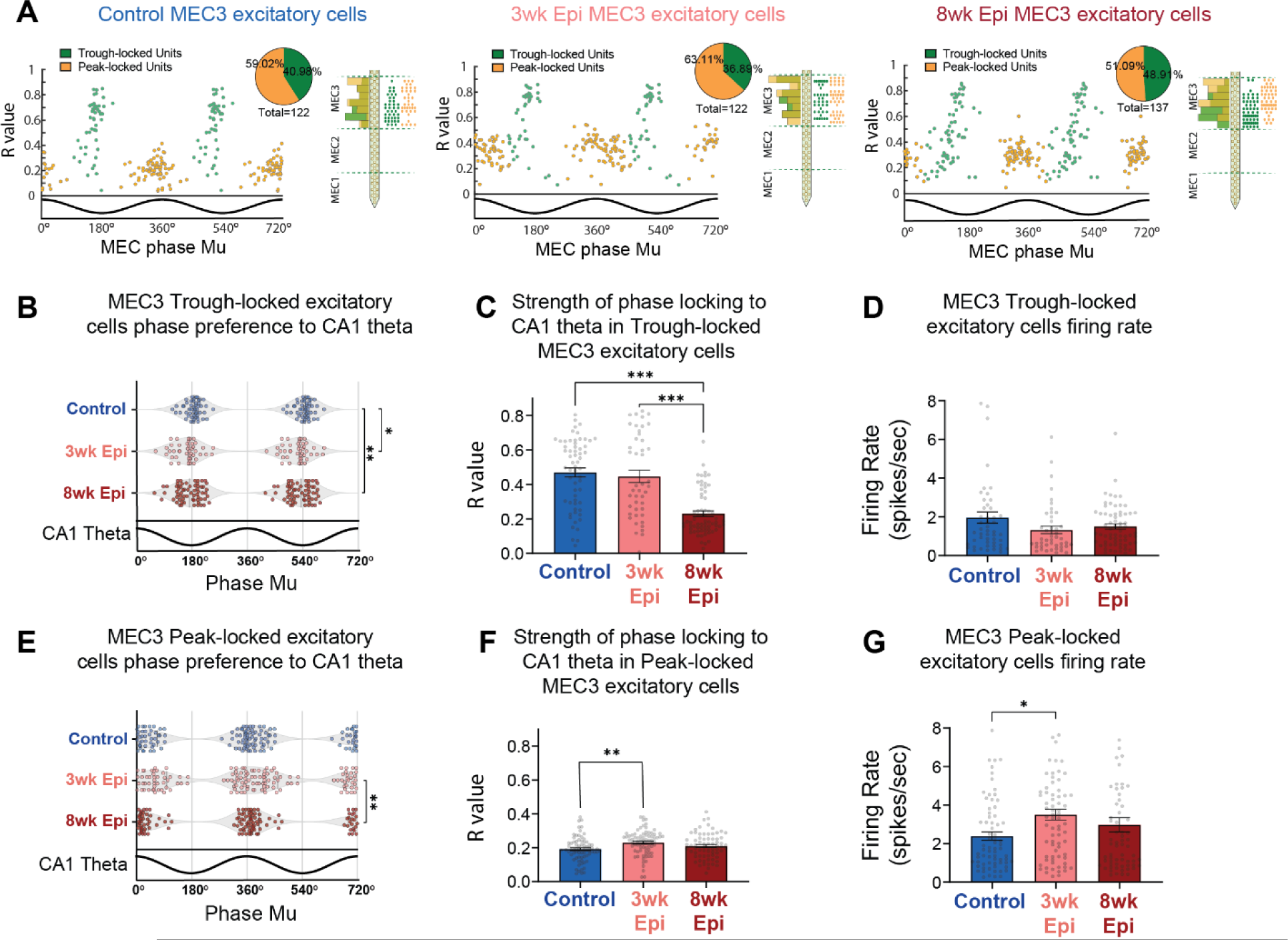
Reduced theta phase locking is specific to a subpopulation of Trough-locked MEC3 excitatory units. **A.** MEC3 excitatory cells show two distinct clusters based on their theta phase preference to local theta (x axis) and phase locking strength (y axis). K-means clustering was used to separate the two populations into Trough-locked units (green) and Peak-locked units (orange). Right side plots show the distribution of the 2 clusters along the probe in each group. **B.** Altered distribution of phase preferences to CA1 theta for MEC3 Trough-locked excitatory units in Epileptic mice. Each data point represents one single-unit and data is double plotted for visualization. (n = 50, Control cells; n = 44, 3wk Epileptic cells; n = 65 8wk Epileptic cells; circular k-test for equal concentrations: Control v. 3wk Epileptic *p* < 0.017, Control v. 8wk Epileptic *p* < 0.003; post-hoc correction for group comparisons by using alpha = 0.017). **C.** Reduced phase locking strength of MEC3 Trough-locked excitatory units to CA1 theta in 8wk Epileptic mice (n = 50, Control cells; n = 45, 3wk Epileptic cells; n = 67, 8wk Epileptic cells; one-way ANOVA, *F* (2, 159) = 39.3, *p* < 0.0001; Bonferroni corrected: Control v. 8wk Epileptic *p* < 0.0001, 3wk Epileptic v. 8wk Epileptic *p* < 0.0001). **D.** No changes in firing rate of MEC3 Trough-locked excitatory units in Epileptic mice (n = 50, Control cells; n = 45, 3wk Epileptic cells; n = 67, 8wk Epileptic cells; one-way ANOVA, *F* (2, 159) = 2.3, *p* > 0.05). **E.** Phase preference to CA1 theta for MEC3 Peak-locked excitatory units in Control and Epileptic animals. Each data point represents one single-unit and data is double plotted for visualization. (n = 71, Control cells; n = 76, 3wk Epileptic cells; n = 70 8wk Epileptic cells; circular k-test for equal concentrations: 3wk Epileptic v. 8wk Epileptic *p* < 0.003; post-hoc correction for group comparisons by using alpha = 0.017). **F.** Increased phase locking strength of MEC3 Peak-locked excitatory cells to CA1 theta in Epileptic mice 3 weeks after Pilo-SE (n = 72, Control cells; n = 77, 3wk Epileptic cells; n = 70, 8wk Epileptic cells; one-way ANOVA, *F* (2, 216) = 5.1, *p* < 0.01; Bonferroni corrected: Control v. 3wk Epileptic *p* < 0.01). **G.** Increased firing rate of MEC3 peak-locked excitatory cells in Epileptic mice 3 weeks after Pilo-SE (n = 72, Control cells; n = 77, 3wk Epileptic cells; n = 70, 8wk Epileptic cells; one-way ANOVA, *F* (2, 216) = 4.2, *p* < 0.05; Bonferroni corrected: Control v. 3wk Epileptic *p* < 0.05). Error bars represent s.e.m. *p<0.05, **p<0.01, ***p<0.001 for C, D, F, G; *p<0.017, **p<0.003 for B, E

We then investigated how each subpopulation of MEC3 excitatory neurons was phase locked to downstream CA1 theta (note that clustering was performed on local MEC theta), and found that the Trough-locked MEC3 neurons were specifically altered in Epileptic mice (Figure 5) and fully accounted for the reduced phase locking strength observed in the full population (Figure 4E). In Epileptic mice, Trough-locked cells showed more dispersed phase preferences to CA1 theta in both 3wk and 8wk Epileptic groups (Figure 5B). However, the strength of theta phase locking (r) was substantially reduced only in the 8-week Epileptic group (Figure 5C). These changes were not driven by changes in overall activity as we found no differences in the firing rate of Trough-locked neurons (Figure 5D). In the Peak-locked cluster, the distribution of preferred phases was similar between the Control group and each Epileptic group (Figure 5E). In addition, there was a small, but statistically significant, increase in phase locking strength to CA1 theta (Figure 5F) as well as in the firing rate (Figure 5G) of MEC3 Peak-locked excitatory cells in 3-week Epileptic animals. Notably, these opposing results in MEC3 Peak-locked and Trough-locked excitatory cells suggests the observed effects are not due to changes in the downstream CA1 theta oscillation, and rather reflected specific changes to the timing of these distinct MEC3 cell types. Together, these results indicate that the late-onset deficits in the strength of MEC3 excitatory cell phase locking in Epileptic mice (Figure 4) are driven entirely by the Trough-locked subcluster. While MEC3 excitatory cells are traditionally treated as a homogeneous population, these results suggest that a unique subpopulation may be specifically vulnerable during the progression of epilepsy and highlight the need for further investigation into these cell types.

### Progressive deficits in theta coherence within MEC and between MEC-HPC emerge between 3 and 8 weeks after Pilo-SE

Both HPC and MEC are required for successful spatial memory performance, and theta synchrony within and between these regions is key to proper spatial processing^28–30,81,82^. Therefore, we examined theta coherence within MEC and between MEC and HPC across the progression of memory deficits from 3 to 8 weeks after Pilo-SE. We found that theta coherence between MEC2 and MEC3 was fully intact at 3 weeks after Pilo-SE, but was reduced by 8 weeks after Pilo-SE (Figure 6A). Similarly, we also found reduced theta coherence between MEC and HPC at 8 weeks after Pilo-SE that was not present at the 3-week time point (Figure 6B). Specifically, by 8 weeks after Pilo-SE, we found reduced theta coherence between MEC1/MEC2 and stratum oriens (Or), stratum pyramidale (Pyr), lacunosum moleculare (LM), and molecular layers (Mol) of HPC (Figure 6B). Notably, these deficits in long-range coherence were restricted to MEC1 and MEC2, with no significant changes in MEC3 coherence with HPC. These changes may reflect altered timing of inputs into superficial MEC layers, and could in turn contribute to the disrupted timing we observed in Trough-locked MEC3 neurons. Together, these data suggest that deficits in theta synchrony within MEC and between MEC and HPC emerge between 3 and 8 weeks after Pilo-SE, during the period that spatial memory impairments are becoming more severe (Figure 1C-D).

**Figure 6:**
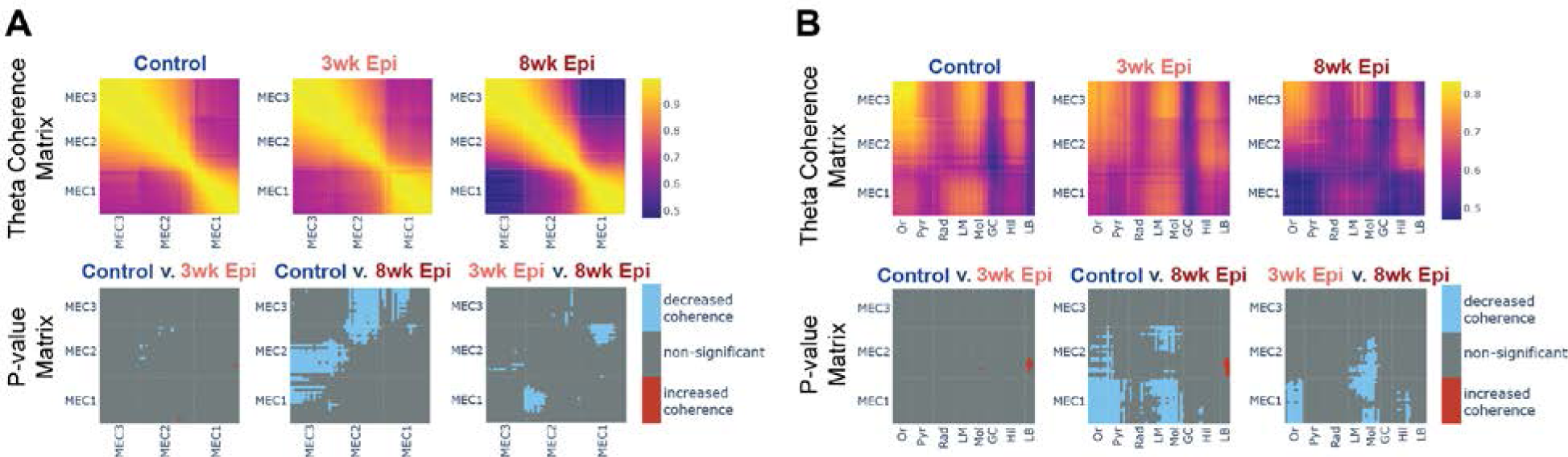
Decreased theta coherence within MEC and between MEC-HPC emerges between 3 and 8 weeks after Pilo-SE. **A.** Theta coherence between each channel pair along the probe in MEC in Control (top left), 3wk Epileptic (top middle) and 8wk Epileptic (top right) mice. P value matrix (bottom row) shows significant changes in coherence at each location between experimental groups, with reduced coherence emerging by 8 weeks after Pilo-SE (N = 11, Control animals; N = 11, 3wk Epileptic animals; N = 9, 8wk Epileptic animals; welch t-test with post-hoc correction by using alpha = 0.017; blue: decrease coherence; red: increase coherence). **B**. Theta coherence between each channel pair along the probe in MEC and HPC in Control (top left), 3wk Epileptic (top middle) and 8wk Epileptic (top right) mice. P value matrix (bottom row) shows significant changes in coherence at each location between experimental groups, with reduced coherence emerging by 8 weeks after Pilo-SE (N = 11, Control animals; N = 11, 3wk Epileptic animals; N = 9, 8wk Epileptic animals; welch t-test with post-hoc correction by using alpha = 0.017; blue: decrease; red: increase). Or, stratum oriens; Pyr, stratum pyramidale; Rad, stratum radiatum; LM, stratum lacunosum moleculare; Mol, molecular layer of dentate gyrus; GC, granule cell layer; Hil, hilus of dentate gyrus; LB, lower blade of granule cell layer.

To test whether these theta coherence deficits in MEC and between MEC and HPC were related to seizure susceptibility, we examined the correlation between theta coherence and seizure frequency and seizure recency. We found that the coherence within MEC or between MEC-HPC was uncorrelated with both seizure frequency or seizure recency (Figure S5E-H), suggesting that these theta coherence deficits are not directly associated with seizure susceptibility.

### Dissociation of long-range MEC-HPC theta coherence from local HPC or MEC coherence deficits

In our recordings, we found several synchronization deficits in Epileptic mice, but little is known about how these deficits may impact each other. To investigate the relationship between local deficits in theta coherence within HPC or MEC and long-range deficits between the two regions, we used a subsampling approach to determine whether these effects could be dissociated. In the 3wk and 8wk Epileptic animals, we selected a subset of time bins that had MEC-HPC theta coherence levels equivalent to Controls. This led to a new “Subsampled-MEC/HPC” dataset with similar MEC-HPC coherence across all groups (Figure 7A, S7, see Methods). We then tested whether theta coherence within MEC or within HPC was still reduced in the Epileptic animals compared to Controls in this subsampled dataset. We found that theta coherence deficits within HPC and within MEC persisted in Epileptic animals in the Subsampled-MEC/HPC dataset (Figure 7A), despite similar cross-region coherence to Controls during the subsampled periods. This indicates that long-range theta synchrony between MEC-HPC can be dissociated from within-region coherence and that normalizing long-range coherence is not sufficient to normalize theta coherence within HPC or within MEC. Furthermore, this suggests that the deficits in long-range (MEC-HPC) and local (within MEC and HPC) coherence that we observed in Epileptic mice are likely driven by distinct circuit mechanisms.

**Figure 7:**
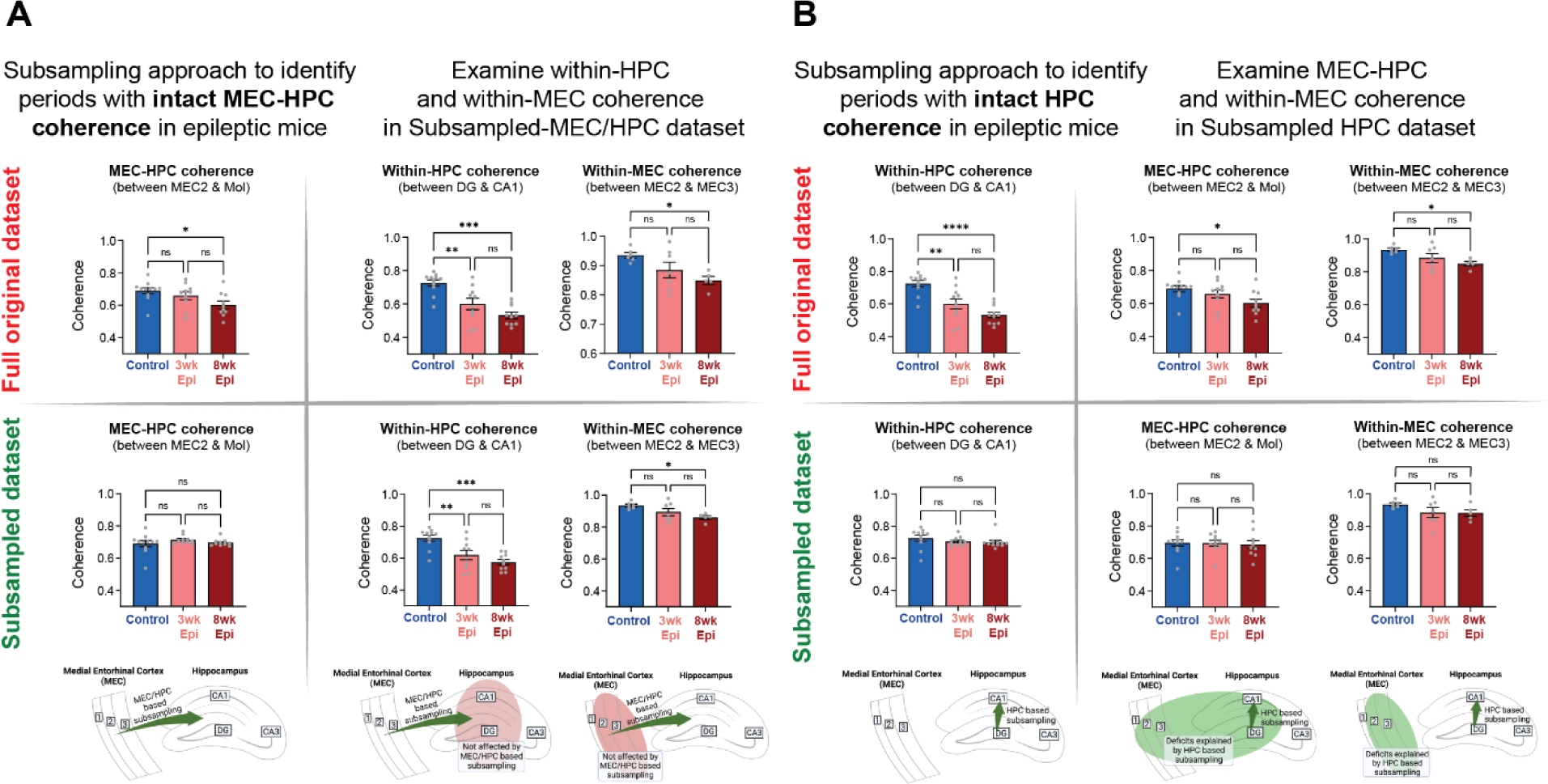
Subsampling approach reveals dissociation between local and long-range theta coherence deficits in Epileptic mice as well as periods of intact theta coherence across the HPC-MEC circuit. **A.** Using a subsampling approach, we identified a set of time bins with equivalent theta coherence between MEC-HPC in all groups showing in left column (original dataset (top left): N = 11, Control; N = 10, 3wk Epileptic; N = 10, 8wk Epileptic; one-way ANOVA, *F(2,28)* = 4, *p* < 0.05, post hoc tests corrected by Bonferroni: Control v. 8wk Epileptic *p* < 0.05, subsampled dataset (bottom left): N = 11, Control; N = 10, 3wk Epileptic; N = 10, 8wk Epileptic; one-way ANOVA, *F(2,28)* = 0.89, *p* > 0.05). Using the full original dataset, theta coherence was reduced within-HPC (top middle, N = 11, Control; N = 11, 3wk Epileptic; N = 11, 8wk Epileptic; one-way ANOVA, *F(2,28)* = 16.1, *p* < 0.0001, post hoc tests corrected by Bonferroni: Control v. 3wk Epileptic *p* < 0.01, Control v. 8wk Epileptic *p* < 0.0001) and within-MEC in Epileptic mice (top right, N = 6, Control; N = 7, 3wk Epileptic; N = 5, 8wk Epileptic; one-way ANOVA, *F(2,15)* = 4.4, *p* < 0.05, post hoc tests corrected by Bonferroni: Control v. 8wk Epileptic *p* < 0.05). In the Subsampled-MEC/HPC dataset, with no deficits in MEC-HPC coherence, within-HPC and within-MEC theta coherence deficits were still present in Epileptic mice (bottom middle, within-HPC: N = 11, Control; N = 10, 3wk Epileptic; N = 10, 8wk Epileptic; one-way ANOVA, F(2, 28) = 12.7, *p* = 0.0001; post hoc tests corrected by Bonferroni: Control v. 3wk Epileptic *p* < 0.01; Control v. 8wk Epileptic *p* < 0.0001; bottom right, within-MEC: N = 6, Control; N = 7, 3wk Epileptic; N = 5, 8wk Epileptic; one-way ANOVA, F(2, 15) = 4.4, *p* < 0.05; post hoc tests corrected by Bonferroni: Control v. 8wk Epileptic *p* < 0.05). **B.** Using a similar subsampling approach, we identified a new set of time bins with equivalent theta coherence within-HPC in all groups showing in left column (original dataset (top left): N = 11, Control; N = 11, 3wk Epileptic; N = 11, 8wk Epileptic; one-way ANOVA, *F(2,30)* = 17.8, *p* < 0.0001, post hoc tests corrected by Bonferroni: Control v. 3wk Epileptic *p* < 0.01; Control v. 8wk Epileptic *p* < 0.0001; subsampled dataset (bottom left): N = 11, Control; N = 11, 3wk Epileptic; N = 11, 8wk Epileptic; one-way ANOVA, *F(2,30)* = 1, *p* > 0.05). In the full original dataset, theta coherence was reduced between MEC-HPC (top middle, N = 11, Control; N = 10, 3wk Epileptic; N = 10, 8wk Epileptic; one-way ANOVA, *F(2,28)* = 4, *p* < 0.05, post hoc tests corrected by Bonferroni: Control v. 8wk Epileptic *p* < 0.05) and within-MEC (top right, N = 6, Control; N = 7, 3wk Epileptic; N = 5, 8wk Epileptic; one-way ANOVA, *F(2,15)* = 4.4, *p* < 0.05, post hoc tests corrected by Bonferroni: Control v. 8wk Epileptic *p* < 0.05). In the subsampled-HPC dataset, with no deficits in within-HPC theta coherences across groups, there were no deficits in long-range MEC-HPC coherence in Epileptic mice (bottom middle, N = 11, Control; N = 10, 3wk Epileptic; N = 10, 8wk Epileptic; one-way ANOVA, F(2, 28) = 0.1, *p* > 0.05), nor any deficits in within-MEC theta coherence in Epileptic mice (bottom right, N = 6, Control; N = 7, 3wk Epileptic; N = 5, 8wk Epileptic; one-way ANOVA, F(2, 15) = 1.8, *p* > 0.05). Error bars represent s.e.m. *p<0.05, **p<0.01, ***p<0.001

Similarly, we next created a second subset of time bins in the Epileptic animals that had levels of theta coherence within HPC (between DG and CA1) equivalent to Controls. This led to a “Subsampled-HPC” dataset that had similar within-HPC coherence across all groups (Figure 7B). We then tested whether Epileptic mice continued to have reduced coherence within MEC or between MEC-HPC in this dataset. Surprisingly, we found that Epileptic mice at both 3 and 8 weeks after Pilo-SE had no deficits in MEC or MEC-HPC theta coherence within this subsample (Figure 7B). That is, in Epileptic mice, periods of normal theta coherence within the HPC were associated with normal levels of MEC and MEC-HPC theta coherence. Furthermore, we found relatively high overlap in time bins between the Subsample-HPC and the Subsample-HPC/MEC and Subsample-MEC datasets (Figure S7B). This suggests there is an intact theta generator in Epileptic mice, even at 8 weeks after Pilo-SE, that can drive coherent theta within and between these regions, but does so only in a subset of time bins. This network-wide theta synchronization may reflect intact long-range synchronization from other brain regions (such as the medial septum or supramammillary nucleus) that are active during specific brain states. Alternatively, it is possible that intact synchronous HPC activity may feed back into MEC to align the activity of these regions in time. Regardless, these results demonstrate that Epileptic mice are capable of synchronizing the MEC-HPC circuit during a subset of time bins.

## DISCUSSION

Together, our work demonstrates that there is a progression of HPC and MEC circuit deficits that arise following epileptogenesis and accumulate as spatial memory impairment worsens. We found that deficits in hippocampal theta and interneuron spike timing emerged early, within 3 weeks after Pilo-SE (Figure 3), while deficits in MEC spike timing, within-MEC theta coherence, and coherence between MEC and HPC emerged later, by 8 weeks after Pilo-SE (Figure 4-6). In particular, a distinct subpopulation of Trough-locked MEC layer 3 excitatory neurons was specifically impaired in Epileptic mice. This progression of theta synchronization and spike timing deficits occurred alongside progressive deficits in spatial memory (Figure 1C-D). Notably, seizures and cell death occurred early after Pilo-SE and did not increase across the development of progressive memory deficits (Figure 2B, Figure S2). Finally, we found that changes in long-range MEC-HPC theta coherence are dissociable from within-region theta coherence deficits in Epileptic mice (Figure 7A), and that there were periods of time when theta coherence within and between all regions was fully intact in Epileptic mice (Figure 7B). Together, these results indicate that 1) in Epileptic mice, neural synchrony within and between HPC and MEC breaks down at distinct time points during the progression of chronic TLE, 2) early changes in HPC synchrony coincide with more subtle memory impairment, while late-onset changes in MEC and between MEC-HPC match the time course of more severe, progressive cognitive deficits, 3) a distinct subpopulation of MEC3 Trough-locked excitatory neurons may be uniquely vulnerable during the progression of epilepsy, and 4) there are multiple distinct drivers of theta in HPC and MEC circuits, with deficits across these circuits occurring at dissociable time points within a session and across the development of epilepsy.

### Progressive memory deficits in Epileptic mice are dissociable from seizures and cell death

Memory deficits are well-established in epilepsy, but little is known about how they progress throughout epilepsy and which specific circuits are involved in producing these memory deficits. Our previous work suggested that spatial representations in CA1 were impaired after Pilo-SE, but there was a dissociation between disrupted spatial precision, which emerged within 3 weeks, and reduced spatial stability, which emerged 6-8 weeks after Pilo-SE^36^. We therefore hypothesized that the expression of memory deficits may also have distinct time courses. Indeed, we found that Epileptic mice had both mild, early-onset memory deficits that progressively worsened after Pilo-SE (Figure 1). These results reinforce the notion that there is an accumulated breakdown of spatial memory circuits in Epileptic mice that contribute to altered memory performance. Meanwhile, seizure frequency and cell death did not progress from 3 to 8 weeks after Pilo-SE (Figure 2B, S2), suggesting that, while they may contribute to the mild, early memory deficits that were present at 3 weeks after Pilo-SE, they are unlikely to drive the progression or worsening of these memory deficits later on. However, it remains possible that the cumulative number of seizures, rather than the onset or frequency, could impact memory performance. Future work using novel approaches to inhibit seizures in this model could examine whether cumulative seizure activity directly contributes to the progression of memory impairment between 3 and 8 weeks after Pilo-SE. Similarly, it remains possible that the indirect effects of cell death (e.g., compensatory excitability changes, axonal sprouting, altered connectivity) may still be progressive and contribute to progressive memory impairment in epilepsy. In addition, because our work was limited to the novel object location memory task, future work will be necessary to further tease apart which specific spatial memory domains are impacted after Pilo-SE.

### Hippocampus is an early site of abnormal synchronization in Epileptic mice

After finding distinct deficits in spatial memory (Figure 1) and spatial coding^36^ at 3 and 8 weeks after Pilo-SE, we next characterized how neural synchronization, and specifically theta, was altered at these time points using *in vivo* electrophysiology. At the 3-week time point, we found that theta synchronization deficits in Epileptic mice were predominantly restricted to the hippocampus. In particular, we found early deficits in HPC theta power, within-HPC theta coherence, and altered theta phase locking of DG interneurons in Epileptic mice (Figure 3). Notably, these changes in HPC were independent of upstream changes in MEC, as we found MEC theta oscillations and spike timing were predominantly intact 3 weeks after Pilo-SE (Figure 4-6). In addition, we found that HPC changes in theta coherence cannot be explained by MEC-HPC coherence deficits (Figure 7A). Together, these results suggest that HPC activity is selectively altered in the first 3 weeks after Pilo-SE, and these deficits are not driven by abnormal inputs from MEC. This time course of HPC desynchronization aligns with poor spatial memory on the Hard NOL task (Figure 1) and reduced precision of spatial coding in CA1^36^. This suggests that disrupted within-HPC synchronization may contribute to the early memory deficits observed at this time point. Notably, even after the onset of major synchronization deficits, we found that during periods with high within-HPC theta coherence, there were also high levels of synchronization in MEC and between MEC-HPC (Figure 7B). This suggests that HPC coherence may be a reliable biomarker for periods of “normal” cognitive function, and could be used to inform neurofeedback interventions or stimulation protocols for responsive neurostimulation. In addition, this suggests that an intact theta generator (possibly long-range inputs from outside HPC and MEC) could be therapeutically targeted or strengthened to restore theta across both the HPC and MEC after epileptogenesis.

### Synchronization deficits in MEC emerge as memory impairment progresses in epileptic mice

We found several deficits in neural synchrony within the MEC and between MEC-HPC, and these changes predominantly emerged between 3 and 8 weeks after Pilo-SE (Figure 4-6). This suggests that MEC circuits are progressively altered following Pilo-SE, while memory deficits progress on the Easy NOL task (Figure 1C-D) and impaired CA1 spatial stability^36^ emerges. This disrupted synchrony may specifically drive changes in hippocampal spatial coding and memory, as proper timing of inputs is required for inducing plasticity mechanisms. For instance, behavioral timescale synaptic plasticity (BTSP) is sufficient to form spatial maps and relies on temporally precise convergent inputs from MEC3 and CA3 to induce new stable place fields in CA1^77–80^. Therefore, in Epileptic mice, the disrupted MEC-HPC theta synchrony may directly impair the development of stable place fields. Notably, this altered MEC synchronization does not coincide with the onset of hippocampal desynchronization, which was present at 3 weeks after Pilo-SE (Figure 3), reinforcing the notion that distinct mechanisms are driving the early- and late-onset circuit deficits in Epileptic mice.

### A theta Trough-locked excitatory population in MEC3 may be selectively vulnerable in epilepsy

MEC3 excitatory cells are typically considered a homogeneous population, but there is evidence of distinct subclasses that are predominantly active at the peak or trough of theta^33^. We found clear evidence of these distinct subpopulations in MEC3 of both Control and Epileptic mice (Figure 5A). Prior work has shown that both Trough-locked and Peak-locked MEC3 neurons contain grid and head direction coding, however the Trough-locked population has more precise tuning (i.e., higher information content and head direction scores)^33^. Surprisingly, we found a subtype-specific deficit in theta phase locking in the Trough-locked MEC3 neurons of Epileptic mice 8 weeks after Pilo-SE (Figure 5C). This deficit was progressive, emerging between 3 and 8 weeks after Pilo-SE, suggesting it may be linked to the progressive memory deficits observed at this time point. Importantly, these progressive deficits do not appear to be driven by cell degeneration in MEC^83–88^, as we found that cell death and FluoroJade C labeling occurred early after Pilo-SE, mostly within the first two days (Figure S2). We also found that reduced NeuN staining was restricted to the ventral portion of MEC3 (Figure S2L-M), while our electrophysiological recordings targeted dorsal MEC.

Notably, several previous studies have found hyperexcitability, specifically in MEC2 circuits in epileptic rats using *ex vivo* brain slices^89–91^. In addition, a recent study has reported that increased excitatory inputs from MEC into the DG may drive chronic seizures in a Dravet Syndrome mouse model^92,93^. However, in the present study, we did not observe increased firing rates during active navigation in MEC2 or MEC3 excitatory neurons using *in vivo* recording in epileptic mice, suggesting that compensatory mechanisms, such as increased local MEC inhibitory inputs, may be sufficient to normalize firing patterns in MEC in the Pilo-SE model.

Furthermore, since there are multiple excitatory cell types in MEC2 and we did not have cell-type specificity in our recordings, it remains possible that specific changes to the HPC-projecting stellate cells could be occluded by changes in other cell types (i.e., MEC2 pyramidal cells). Further work is needed to investigate the post-synaptic excitability of MEC-HPC projection specific neurons. In addition, the selective nature of the phase locking deficit in MEC3 suggests that there is a specific vulnerability of the Trough-locked cells, which could be driven by differences in intrinsic properties or connectivity profiles. Currently, very little is known about this specific subtype and therefore future work is necessary to determine its genetic, anatomical, and functional profile.

One critical outstanding question is whether the precision and stability of MEC spatial representations (i.e., grid, head-direction, border coding) may also be impaired at these time points. While our recordings were not designed to investigate changes in MEC spatial coding, future studies with chronic freely behaving recordings will be necessary to determine how the spatial content of inputs to HPC are altered in epileptic mice.

### Dissociable circuit mechanisms drive within- and cross-region theta coherence deficits

Our work has identified several synchronization deficits in epileptic mice within and between HPC and MEC. To examine how these deficits might be related, we performed a subsampling analysis to determine if periods of normal (i.e., equal to Controls) cross-region theta coherence (MEC-HPC) were associated with normal within-region coherence (Figure 7A, S7). If within- and cross-region coherence deficits were controlled by a single network mechanism (e.g., collateral long-range inputs), then they should be correlated in time throughout the recording. However, we found a clear dissociation between these deficits in Epileptic mice, suggesting that distinct networks were contributing to local and long-range desynchrony. This suggests that MEC-HPC coherence is not sufficient for downstream HPC coherence and that these within- and across-regions deficits can be dissociated. On the other hand, when we subsampled periods of normal within-HPC theta coherence, we found that these periods were associated with normal levels of MEC and MEC-HPC coherence (Figure 7B). This suggests that there is a theta generation mechanism that can influence both the HPC and MEC that is relatively intact in epileptic mice, but only drives intact entorhinal-hippocampal synchronization in a subset of time periods.

Notably, using this subsampling approach, we were able to find periods of intact theta coherence between HPC and MEC (Figure 7A), within-HPC (Figure 7B), and within-MEC (Figure S7B). We further explored how these states related to each other by examining the proportion of time bins that overlapped within each of these subsampled datasets. Consistent with our results on the relationship between theta coherence in MEC and HPC (Figure 7), we found relatively high overlap across each of our subsampled datasets (Figure S7B), and particularly for the within-HPC subsample. Together, these findings suggest that theta synchronization within and between MEC and HPC remains structured in Epileptic mice and that periods of intact HPC theta coherence represent time bins where coherence across the MEC-HPC circuit is highly coordinated (Figure S7C). There are several possible candidate mechanisms that could drive this intact theta in Epileptic mice, including long-range projections from the medial septum or supramammillary nucleus, or feedback projections from CA1 to MEC. Future work is needed to determine how each of these projections are impacted by epilepsy and whether they can control entorhinal-hippocampal synchronization in epileptic animals.

### New technical developments are needed to directly test the causal role of altered synchronization in epilepsy

The recording techniques utilized here are ideal for examining how specific neural circuits are altered during the progression of memory impairments in epilepsy. But while altered theta synchronization is likely to impact information processing in the hippocampal-entorhinal system, it is difficult to draw causal conclusions from recordings alone. Therefore, it is critical that the field develop new methods to directly manipulate synchronization in awake, behaving animals to determine the precise role of theta synchrony in behavior. New experimental tools that deliver electrical or optogenetic stimulation in phase with endogenous rhythms^94,95^ can be used to determine how the timing of neural activity directly impacts behavior. These new methods of stimulation can lead to increased coherence between regions^95^ or change the theta phase locking in distinct subpopulations of neurons^94^. Our work here suggests that resynchronizing the timing of DG inhibitory neurons and trough-locked MEC3 neurons, or increasing theta coherence between MEC and HPC may lead to improved memory in epileptic mice. Future work will be necessary to directly test the causal role of each of the theta synchronization deficits that we have identified in epileptic mice.

## STAR METHODS

### KEY RESOURCES TABLE

| REAGENT or RESOURCE | SOURCE | IDENTIFIER |
| --- | --- | --- |
| **Antibodies** |  |  |
| Rabbit anti-NeuN | Millipore Sigma | Cat# AB78N; RRID: AB_10807945 |
| FluoroJade C | Biosensis | Cat# TR-100-FJ |
| Alexa Goat Anti-Rabbit 555 | Thermo Fisher Scientific | Cat# A21429; RRID:AB_2535850 |
| Alexa Goat Anti-Rabbit 488 | Thermo Fisher Scientific | Cat# A11008; RRID:AB_143165 |
| **Chemicals, peptides, and recombinant proteins** |  |  |
| Pilocarpine Hydrochloride | Sigma-Aldrich | Cat# P6503; CAS# 54-71-7 |
| Scopolamine Methyl Nitrate | Sigma-Aldrich | Cat# S2250 ; CAS# 155-41-9 |
| Diazepam | DASH Pharmaceuticals | NDC# 69339-137-01 |
| DPX | Thermo Fisher Scientific | Cat# 50980369 |
| Histo Clear II (Xylene) | Thermo Fisher Scientific | Cat# 5032947 |
| DAPI | Thermo Fisher Scientific | Cat# D1306 |
| Trypsin | gibco | Cat# 15090-046 |
| Dil Dye | Invitrogen | Cat# V22885 |
| Isoflurane | Baxter Healthcare Corporation | NDC# 10019-360-40 |
| Paraformaldehyde | Wako | Cat# 163-20145 |
| HEPES | Sigma-Aldrich | Cat# V900477 |
| NaCl | Sigma-Aldrich | Cat# V900058 |
| KCl | Sigma-Aldrich | Cat# V900068 |
| CaCl<sub>2</sub> | Sigma-Aldrich | Cat# V900269 |
| MgCl<sub>2</sub> | Sigma-Aldrich | Cat# V900020 |
| Triton X-100 | Thermo Fisher Scientific, Fisher BioReagents | Cat# BP151-100; CAS# 9002-93-1 |
| Normal Goat Serum | Calbiochem | Cat# NS02L |
| phosphate buffered saline (PBS) | Thermo Fisher Scientific | Cat# bp339-500 |
| Carprofen | Rimadyl (Through Mount Sinai Pharmacy) | Cat# 4019449 |
| Lidocaine (Xylocaine 2%) | Premier ProRX (Through Mount Sinai Pharmacy) | Cat# PRX480627 |
| Ampicillin | Sigma Aldrich | Cat# A0166-5G |
| Experimental Models: Organisms/strains |  |  |
| Mouse: C56BL/6 | Charles River | Strain# 027 |
| Mouse: 129X1/SvJ (OVX mice) | Jackson Lab | Strain# 000691; RRID:IMSR_JAX:000691 |
| Software and algorithms |  |  |
| MATLAB | The Mathworks, Inc | https://www.mathworks.com/ |
| Python | Open-source | https://www.python.org/ |
| Prism 9.3 | GraphPad Software Inc | https://www.graphpad.com/updates/prism-931-release-notes |
| NeuroScore | DSI | Version 3.3.9318 |
| Biorender | Biorender | https://www.biorender.com/ |
| ImageJ | NIH | https://imagej.net/Fiji |
| Adobe Illustrator | Adobe | n/a |
| Miniscope recording software | Open-source | https://github.com/Aharoni-Lab/Miniscope-DAQ-QT-Software |
| Chronotate | Open-source | https://github.com/ShumanLab/Chronotate |
| Cell counter | Open-source | https://github.com/ZachPenn/CellCounting |

### RESOURCE AVAILABILITY

#### Lead Contact

Further information and requests for resources or raw data should be directed to and will be fulfilled by the Lead Contact, Tristan Shuman.

### EXPERIMENTAL MODEL AND SUBJECT DETAILS

#### Study Design

All experimental procedures were approved by the Icahn School of Medicine’s Institutional Animal Care and Use Committee (IACUC), in accordance with the US National Institutes of Health (NIH) guidelines. For EEG recordings, we used wireless EEG devices from Data Science International (DSI) to continuously monitor seizure activity prior to silicon probe recordings. For silicon probe acute recordings, we used two 256-channel probes from UCLA Masmanidis Lab to simultaneously record in HPC and MEC at 3 or 8 weeks after pilocarpine-induced status epilepticus (Pilo-SE).

#### Animals

Adult male C57BL/6 mice (Charles Rivers Laboratories) were used in all experiments. Male mice had lower mortality rates in preliminary studies and were previously utilized to identify changes in the progression of spatial coding deficits after Pilo-SE^36^. We therefore restricted our sample to only male mice in this study. Outside of water restriction, mice were given ad libitum food and water on a 12-hour light-dark cycle. Mice were group housed with 4-5 per cage until wireless EEG recording started. Mice were then separated into individual housing for recording purposes and co-housed with one ovariectomized female mouse (129X1/SvJ, The Jackson Laboratory). Pilocarpine-induced status epilepticus occurred at 9 weeks of age.

### METHOD DETAILS

#### Wireless EEG and Headbar Implant Surgical Procedures

3-4 weeks before *in vivo* electrophysiology recordings, all mice underwent headbar and wireless EEG implantation surgeries. During the surgery, animals were head fixed on the surgical stereotax (KOPF Instruments) and maintained under anesthesia with 1-3% isoflurane (Baxter Healthcare Corporation, NDC 10019-360-40). The scalp was cleared of hair and sterilized with betadine (Purdue Frederick, NDC 0034-2200-80) and 70% ethanol. Once the skull was exposed, the fascia was cleared with 3% hydrogen peroxide (MEDLINE, NDC: 53329-981-06). The animal’s skull was then aligned to the stereotax using bregma and lambda. To implant the EEG device (Data Science International, model ETA-F10), we enlarged the scalp opening to the neck area and used blunt, bent forceps to open the cavity under the back skin. We then inserted the body of the EEG device into the back underneath the back skin, with two lead wires exposed from the neck area. Two burr holes were then made on different hemispheres. 1mm length of insulation was stripped off from the tip of both wire leads before inserting into the burr hole and securing between the dura and skull with cyanoacrylate glue. To allow for head-fixation during recordings, we secured a stainless steel headbar to the skull using glue and dental cement (LANG Dental Mfg, REF 1330, REF 1306), and then built up the dental cement to create a well around the exposed skull. Kwik-Sil (World Precision Instruments, Cat# 600022) was then used to fill the well. A final layer of cement was then applied on top of the Kwik-Sil and the headbar. Lidocaine (Premier ProRX, Cat# PRX480627, 2%, ∼0.05ml) was injected subcutaneously into the neck area to assist with recovery from EEG implantation. In addition, carprofen (Rimadyl, Cat# 4019449, 5 mg/kg) was administered during surgery and for 2 days after and ampicillin (Sigma Aldrich, Cat# A0166-5G, 20 mg/kg) was administered during surgery and for 6 additional days.

#### Pilocarpine-induced Status Epilepticus

Mice were randomly assigned to receive either saline or pilocarpine injections at 9 weeks of age. On the day of injection, all animals first received a 0.1 ml intraperitoneal injection of scopolamine methyl bromide (0.2 mg/ml, Sigma-Aldrich, Cat# P6503; CAS# 54-71-7) to reduce the peripheral effects of pilocarpine. 30 minutes later, all pilocarpine-assigned mice received intraperitoneal injections of pilocarpine hydrochloride (Sigma-Aldrich, Cat# S2250; CAS# 155-41-9), with the dose depending on their body weight (250 mg/kg if more than 25 g; 275 mg/ml if between 20 and 25 g; 285 mg/kg if less than 20 g). Saline-assigned mice received equivalent volumes of saline (Fresenius Medical Care, Cat# 060-10109) instead. If pilocarpine-treated mice had not seized within 45 minutes of the initial pilocarpine injection, booster injections of 50-100 mg/kg pilocarpine were given. Status epilepticus (SE) was established when animals entered a continuous seizure, and didn’t react to outside stimuli. Pilocarpine-treated mice were left in SE for 2 hours, after which they received a 20 mg/kg intraperitoneal injection of diazepam (DASH Pharmaceuticals, NDC 69339-137-01) to terminate SE. Pilocarpine-treated mice who didn’t reach SE or died during the procedure were excluded from the study (SE success rate ∼75%). All other animals were given 1 ml of saline through subcutaneous injection by the end of the day, access to moistened food, and were monitored for 72 hours to ensure recovery. Additional saline was given if pilocarpine-treated animals were slow to recover. Wireless EEG transmitters were turned on after SE induction in mice recorded 3 weeks after Pilo-SE. For mice recorded 8 weeks after Pilo-SE, EEGs were turned on 5 weeks after Pilo-SE. In all pilocarpine-treated mice, we confirmed spontaneous seizures through EEG recording.

#### Novel Object Location Task

The novel object location task was performed either 3 weeks or 8 weeks after Pilo-SE in both Epileptic and age-matched Control groups. All animals were first habituated to the behavior room for 6 days prior to the test. After being left in the behavior room in their home cage for 20 minutes, mice were handled by the experimenter for 3 minutes each day. Animals were then put in an empty box (1 ft x 1 ft) with different wall patterns on each side for 10 minutes for box habituation on day 7. On the day of the NOL test (day 8), animals were first put into the empty box to explore without objects for 6 minutes. We then took them out, cleaned the box with 70% ethanol, and taped down two identical objects (1.5 inches x 2 inches shiny gold door stoppers) at the two corners of the box. Animals were put back in the box for two 6-minute training sessions of object exploration (for the Hard version), and this 6-minute object exploration period was repeated a third time for the Easy version of the task. The mice were then returned to their home cages for 3-hours before the test session. During the test session, one of the two objects was moved to a novel location, and the animals were allowed to explore the box for 6 minutes. All sessions were recorded with webcams using the v3 Miniscope recording software^36,96^ (available at https://github.com/Aharoni-Lab/Miniscope-DAQ-QT-Software) to record multiple videos simultaneously. Training and test recordings were then scored by a technician who was blinded to the experimental group. Investigation times and durations were recorded using Chronotate^97^ (available at https://github.com/ShumanLab/Chronotate). The animal’s preference to investigate either object was indicated by a discrimination index (DI):

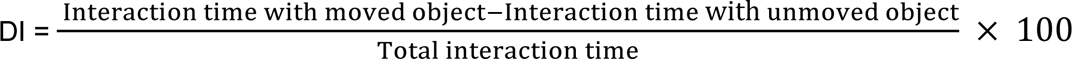

A higher DI during the test means a stronger preference for the moved object, indicating intact spatial memory of the unmoved object. Note that animals with |DI| higher than 30 during training, or that explored either object for less than 3 seconds during training, were excluded from the analysis (N=2 animals excluded).

#### Virtual Reality (VR) Training

Mice were trained to navigate a virtual linear track to obtain water rewards. As such, they were water restricted during training starting from initial handling. Mice had access to 1-2ml of water daily to maintain their body weight around 85% of their initial weight, which was measured right before water deprivation started^36^. More water was offered immediately if mice showed any sign of dehydration or if their body weight was below 78%. Training began with 2 days of handling followed by 2 days of handling plus habituation to head-fixation. Next, mice were introduced to the Styrofoam ball and were given 3 days to habituate to walking on the ball while head-fixed. Then we added lick training, during which animals were trained to lick at a metal tube for water while head-fixed on the ball. Virtual reality (VR) linear track training started after animals could reliably gain water from the lick port. Our VR setup included three 24-inch flat monitor screens (DELL) that angled 120 degrees to each other and surrounded the animal. ViRMEn^98^ (available at http://pni.princeton.edu/pni-software-tools/virmen), an open-source MATLAB software, was used to create the virtual environment. Animals were trained to run down VR tracks of increasing length to release a 5uL water drop at the end of the VR track. Mice were considered ready for silicon probe recording when they could reach at least 100 trials within 1 hour on the longest track (2m in the real world). The VR training phase typically lasted 5-7 days.

#### Craniotomy and Ground Implantation Surgery

Craniotomy and ground wire implantation happened the day prior to acute silicon probe recording. Animals were lightly anesthetized with isoflurane and head-fixed on the stereotax. The top layer of cement was drilled off, and Kwik-Sil was removed to expose the skull. A burr hole was then made over the cerebellum, on the left side. The ground wire (Ag/AgCl-coated, Warner Instruments) was slipped into the burr hole to sit above the dura. A small drop of cyanoacrylate glue was used to secure the ground wire in place. 1.6mm wide craniotomy was then made above HPC (centered at 2mm posterior to bregma, 1.8mm right of bregma) and MEC (centered at 250um anterior to the superior cerebellar artery; 3.3mm right of bregma). The craniotomies were covered with buffered artificial cerebrospinal fluid^36,96^ (ACSF; in mM: 135 NaCl, 5KCl, 5 HEPES, 2.4 CaCl2, 2.1 MgCl2, pH 7.4) and Kwik-Sil was then applied to the skull to cover and protect the craniotomies. Animal received a carprofen (Rimadyl, Cat# 4019449, 5 mg/kg) injection after the craniotomy to reduce pain and prevent inflammation.

#### Acute Silicon Probe Recordings

On the day of recording, we first painted the 256-channel probes (UCLA Masmanidis lab, HPC probe: 4 shanks, 400 um apart; MEC shank: 4 shanks, 200 um apart) with DiI (Invitrogen, Vybrant DiI, V22885) to be able to visualize and check the probe tract after recording. Mice first received carprofen (Rimadyl, Cat# 4019449, 5mg/kg) before being head-fixed onto the Styrofoam ball. We removed the Kwik-Sil and replaced it with buffered ACSF^36,96^. The mouse’s skull was then aligned to the micromanipulator (ROE-200, Sutter Instrument). We then attached the two 256-channel probes to a micromanipulator and lowered them into HPC (coordinates for middle shank: 1.95mm posterior, 1.65mm right, and 2.3mm ventral from bregma) and MEC (coordinates for middle shank: 190um anterior to the superior cerebellar artery, and 3.05mm right, and 3.3mm ventral to bregma) slowly at ∼5um/s. We used a custom MATLAB script to plot raw and filtered data, the theta phase shift, and coherence for each region. We used these electrophysiological features to determine if the probes were in the correct location^34,99–103^. Once probes were both in place, we applied mineral oil (Fisher Chemical, Cat# 8042-47-5) over the buffered ACSF and the brain was allowed to settle for 1 hour.

Electrophysiology recording signal was collected as described previously^104^. Each probe was connected to two 128-channel headstages (Intan Technologies, Cat# C3316) to organize and amplify the signals. Signals were recorded with an Intan recording controller (Intan Technologies, INTAN 1024ch Recording Controller) to collect and log electrophysiology signals from each channel at 25kHz. Licking behavior, reward delivery times, animals’ position on the virtual track, and running speed were logged through the Intan I/O Expander connected to the Intan Recording Controller at the same sampling rate of 25kHz and were saved on a local computer through Intan RHX Data Acquisition Software.

Animals were allowed to run in the 2m long VR track to earn water rewards at least 100 times. Then we switched the animal into a novel environment and allowed another ∼100 trials in the novel context. The average total recording length was ∼180 minutes. Note that all data presented here was taken while animals were traversing the VR track they had been previously trained on (i.e., was familiar).

#### Histology Confirmation of Probe Track

Following the recording, we removed the probes from the animal’s brain and rinsed them with MilliQ water before putting them in 2.5% trypsin (gibco, Cat# 15090-046) for 40 minutes. The mouse was deeply anesthetized using 4% isoflurane before performing a quick decapitation. The brain was extracted and put in 4% paraformaldehyde (PFA; Wako. Cat# 163-20145) and kept at 4°C for 24 hours before changing into fresh phosphate buffered saline (1xPBS; Thermo Fisher Scientific, Cat# bp339-500). The right hemisphere of the brain was sectioned sagittally using the vibratome (Leica VT1000S) at 80 um. Sections were then stained for NeuN and DAPI to help differentiate MEC layers (see Immunohistochemistry). Three excitation wavelengths were used to capture the probe tracts and anatomical structures (DiI, wavelength 555), DAPI (wavelength 408), and NeuN staining (wavelength 488). For each shank, we referenced the mouse brain atlas^105^ to validate the probe location. Only shanks that were in the correct HPC and MEC locations covering all or partial sublayers (CA1-DG layers for HPC; MEC layer 1-3 for MEC) were included in the analysis. Example shanks can be found in Figure S3.

#### Post-processing and analysis of Local Field Potential Data

All data analysis was performed in custom MATLAB and Python scripts. We first took the minute-by-minute raw data and concatenated it into continuous signals for each channel.

For local field potential (LFP) analysis, all data were first down-sampled to 1kHz. We cleaned out 60Hz noise by passing a custom notch filter in MATLAB. Data were then filtered by different bandpass filters with a focus on theta (5-12 Hz) oscillations. Theta phase shift and coherence were then measured using the Chronux toolbox^106,107^. Using electrophysiology features (theta peak power^101^, theta/gamma phase shift^34,100,101^, theta/gamma coherence^99,103^, dentate spike phase reversal^102^, spike density) and probe tract histology images, we were able to identify the location of each channel within the sublayers of HPC and MEC. The power plot (Figure 3A) was generated by averaging theta power for each subregion. Because seizures and locomotion are known to impact theta power^108,109^, we limited our analyses to periods when the animal was locomoting and not seizing (the mouse needed to be moving continuously for more than 3 seconds to be considered locomoting; once a seizure was detected, the seizing period and the 10min after the seizure were excluded from analysis).

Coherence analyses were performed using the Chronux toolbox^106,107^. Each channel pair generated a theta coherence value. To do group analysis, a uniform-shaped sub-region matrix was generated for each animal, and the coherence value for a given region-pair was assigned to the sub-region matrix for group comparison analysis. Coherence data was exported from MATLAB and plotted in Python for visualization (Figure 3B, 6A, 6B). To perform the coherence subsampling analysis (Figure 7), recordings were broken into running bout bins, defined by more than three seconds of continuous locomoting. Then cross region MEC-HPC (MEC2 to DG molecular layer) coherence, within region HPC (DG hilus to CA1 pyramidal layer) coherence, and within region MEC (MEC2 to MEC3 molecular layer) coherence were calculated for each running bin. To determine when Epileptic animals’ theta coherence values were similar to Controls, we selected the time bins in 3-week and 8-week Epileptic animals that showed theta coherence within one standard deviation of the Control group’s coherence value (Figure S7A). Cross-region and within-region coherences were then recalculated within these subsample of time bins (Figure 7A, S7) to study the relationship between coherence deficits of different region pairs. Note that animals were excluded from all subsampled plots if insufficient time bins were available to create a subsampled dataset with coherence equivalent to Controls.

#### Single Unit Spike Sorting and Phase Preference Analysis

To extract single unit activity, the raw electrophysiological data from each shank was first background subtracted in a custom MATLAB script. Background subtracted data was then passed into kilosort 2.5, an automatic spike sorting package using a template-matching approach with drift correction^110,111^. Phy2^110^ was used to manually confirm and clean each unit as needed. All single unit analyses were carried out in MATLAB using custom scripts.

Spike-sorted units were first classified as putative inhibitory and excitatory units in HPC and MEC based on their firing rate, waveform characteristics, and autocorrelogram^34,36,76^. In HPC, units with a firing rate below 8Hz, complex spike index over zero, mean autocorrelogram less than 0.1, and c index (trough-to-peak latency) above 0.26 were classified as excitatory cells. Units with firing rates over 0.2Hz and mean autocorrelogram greater than 0.1 were classified as inhibitory cells (Figure S6A). In MEC, we performed kmeans clustering based on c index (trough-to-peak latency) to separate inhibitory and excitatory units^34^ (Figure S6B).

We quantified phase preference to local or downstream theta oscillation by measuring the mean phase of firing (mu value) and phase locking strength (r value) of each cell. Note that only cells with significant r-values (i.e., were significantly phase-locked; Rayleigh’s test for non-uniformity) were included in the phase preference (mu) analysis (see Quantification And Statistical Analysis). To isolate reference theta for phase preference analysis, the raw data was filtered to the bandwidth of interest (5-12Hz), and a representative channel was picked for each region as reference oscillation (middle channel in MEC2/3 as local MEC theta; middle channel in hilus as DG theta; top channel in pyramidal cell layer as CA1 theta). The theta phase angle was then determined using the Hilbert transformation. MEC3 excitatory cells were separated into trough and peak-locked cells using k-means clustering. Two parameters, r value and mu value, were used during the clustering. The number of cells per animal is listed in Table 1.

**Table 1.**
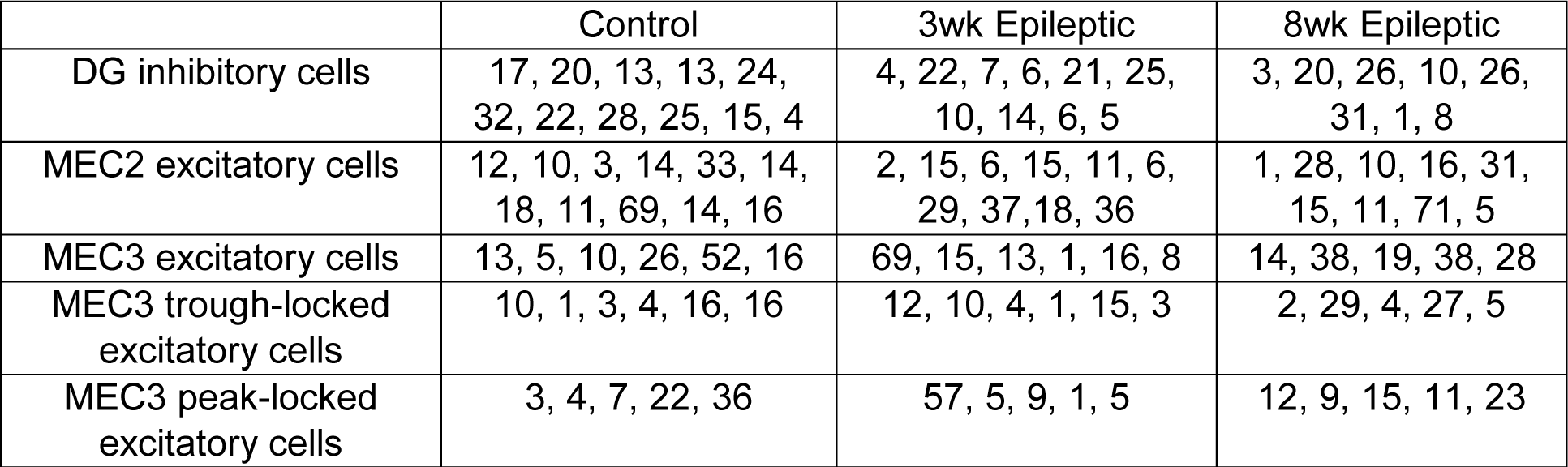
Cells recorded per animal in Control, 3-week Epileptic, and 8-week Epileptic groups.

#### EEG seizure quantification

All EEG seizure detection was performed manually by experimenters who were blinded to the animal group information. Neuroscore (DSI, Version 3.3.9318) was used for EEG signal visualization and seizure detection. EEG seizures were identified by a sharp signal amplitude increase, followed by a 2-10 minute quiet period. EEG signals were not collected when animals were in training or receiving a craniotomy.

#### Immunohistochemistry at Different Timepoints

We used FluoroJade C (FJC) and NeuN staining to characterize cell degeneration and neuron number, respectively. We perfused the animals with 1x PBS (Thermo Fisher Scientific, Cat# bp339-500) and 4% PFA (Wako, Cat#163-20145) at 2 days, 3 weeks, or 8 weeks after status epilepticus. Extracted brains were post-fixed in 4% PFA for 12-24 hours before being transferred into a 30% sucrose solution (in 1x PBS). Once the brain sank to the bottom of the sucrose solution (∼48 hours), we cut the brain down the midline and stored the left and right hemispheres in the −80°C freezer.

For NeuN staining, the right hemispheres were sliced sagittally at 40um on a cryostat (Leica CM3050S). Sections containing HPC or MEC underwent two 15-minute washes in 1 X PBS, followed by 2 hours of blocking at room temperature in 0.3% Triton X-100 (Fisher BioReagents, Cat# BP151-100) and 3% normal goat serum (Calbiochem, Cat# NS02L) in 1xPBS (diluted from 10xPBS). Next, slices were incubated in rabbit anti-NeuN (Millipore Sigma, ABN78) primary antibody (1:2000, in blocking solution) at 4°C overnight (∼15 hours). On the next day, slices underwent three 15-minute 1xPBS washes. They were then incubated in secondary antibody in PBS (1:500 goat anti-rabbit polyclonal Alexafluor 555, Thermo Fisher Scientific, A21429) for 2 hours at room temperature while covered in foil. Slices were washed in 1xPBS three times with DAPI (Thermo Fisher Scientific, D1306, at 1:5000) added during the second wash. All washes and incubations took place on a shaker. We then mounted all slices containing HPC and MEC on Super Plus Microscope Slides (Thermo Fisher, 22-037-246) with ProLong Gold mounting solution (Thermo Fisher Scientific, P36930). Slides were sealed with nail polish before being stored at 4°C.

For FJC staining, the left hemispheres were sliced sagittally at 30um on a cryostat (Leica CM3050S). Biosensis (Ready-to-Dilute) Fluoro-Jade C staining kit (TR-100-FJ) was used for staining. HPC- and MEC-containing sections were then selected and mounted on slides (Thermo Fisher, 22-037-246) and dried at 60°C on a slide warmer for 1 hour to allow good adhesion. Slides were then incubated in a Coplin jar (Thermo Fisher, Polypropylene Coplin Staining Jar, 0181621) with 9 parts of 80% ethanol and 1 part of sodium hydroxide (biosensis Fluoro-Jade C staining kit) for 5 minutes. Slides were transferred to a jar containing 70% ethanol for 2 minutes and then into distilled water for 2 minutes. The endogenous fluorescent background was blocked by incubating in 1 part potassium permanganate (biosensis Fluoro-Jade C staining kit) mixed with 9 parts distilled water for 10 minutes. Slides were then immersed in the FJC staining solution, which contained 1 part of the FJC solution (biosensis Fluoro-Jade C staining kit) and 9 parts of distilled water, for 10 minutes in the dark. Three rounds of distilled water rinses were followed to clean the dye from the slices before the slides were dried on a slide warmer at 60°C for 10 minutes in the dark. Finally, slices were cleared by 3 minutes of immersion in xylene (Thermo Fisher, Histo-Clear II 5032947) before being mounted using DPX (Thermo Fisher, 50980369).

All images were taken by a Leica DM6B fluorescence microscope (Figure S2). Cell counting was performed using an automatic cell counting pipeline (available at https://github.com/ZachPenn/CellCounting) in combination with manual counting with the assistance of ImageJ^112,113^.

## QUANTIFICATION AND STATISTICAL ANALYSIS

All statistics were performed in GraphPad Prism 9.3.1 except for the *p*-value matrices and single-unit phase preference (mu; circular data), which were calculated in Python3 and MATLAB (R2021b), respectively.

For the *p*-value matrices, a Welch’s t-test was used between each channel pair across the group. In Supplementary Figures 4C and 4D, the threshold for significance (alpha) was set to 0.05. In Figures 3B and 6, the alpha was set to 0.017 to correct for multiple comparisons among the three groups. Only comparisons with a *p*-value below alpha were marked in blue (decrease coherence) or red (increase coherence) in the *p*-value matrices.

The Circular Statistic Toolbox^114^ in MATLAB was used to calculate and compare circular data (i.e., mu). We performed the circular Kuiper test to compare the distributions of two populations and the circular k-test to test the equality of phase concentration parameters between groups. Only group differences that showed significantly different distributions (Kuiper test) were additionally tested for changes in concentration (circular k-test). The asterisks on Figure 3C, 4A, and 5B, E represent the significant changes in concentration. The threshold for significance (alpha) was placed at *p < 0.017, **p < 0.003, and ***p < 0.0003 to account for multiple comparisons across groups.

All other statistics were performed in GraphPad Prism. All data are presented as mean +/- SEM, with n representing the number of cells and N representing the number of animals. To compare theta power along hippocampal layers between the three experimental groups, the Repeated-measures mixed-effects model followed by Holm-Sidak post hoc correction was used. One-way or two-way ANOVAs followed by Bonferroni’s post hoc test were used to statistically compare normally distributed data. LFP power and coherence changes were tested for correlation with seizure frequency and seizure recency using a Pearson test and reported with correlation coefficient r value (Figure S5). All statistical tests with post hoc correction were two-tailed with thresholds of significance set at *p < 0.05, **p < 0.01, and ***p < 0.001.

Full details for all statistical tests are reported in the figure legends.

## DATA AND CODE AVAILABILITY

We are currently organizing all LFP and spike data and will make them publicly available in the near future. Any additional information required to re-analyze the data reported in this paper is available from the lead contact upon request.

## AUTHOR CONTRIBUTIONS

Conceptualization, Y.F., Z.CW., and T.S.; Methodology, Y.F., L.P-H., L.M.V. and T.S.; Formal Analysis, Y.F., Z.D., and T.S.; Investigation, Y.F., K.S.D., Z.D., L.P-H., V.P-H., J.S., S.I.L., P.A.P., A.J., C.J.R., and N.N.K.; Writing – Original Draft, Y.F. and T.S.; Writing – Review & Editing, Y.F., Z.CW., Z.T.P., L.M.V., D.J.C., and T.S.; Funding Acquisition, D.J.C. and T.S.; Supervision, Y.F., Z.CW., I.S., D.J.C., and T.S.

## ACKNOWLEDGEMENTS

We thank the entire Cai and Shuman lab groups for the feedback, comments, support, and help on this project. This work was supported by CURE Taking Flight Award (to TS), American Epilepsy Society Junior Investigator Award (to TS), R01 NS116357 (to TS), RF1 AG072497 (to TS), R01 MH120162 (to DJC), DP2 MH122399 (to DJC), R56 MH132959 (to DJC), American Epilepsy Society Predoctoral Research Fellowship (to YF), F31 AG069496 (to LV), F99 NS135813 (to IS) and F32 NS116416 (to ZCW). We also thank BioRender for creating an easy platform for figure creation. Parts of Figures 2, 5, 7, and Supplementary Figure 7 were created with BioRender.com.

## DECLARATION OF INTERESTS

The authors declare no competing interests.

## SUPPLEMENTARY FIGURES

**Supplementary Figure 1:**
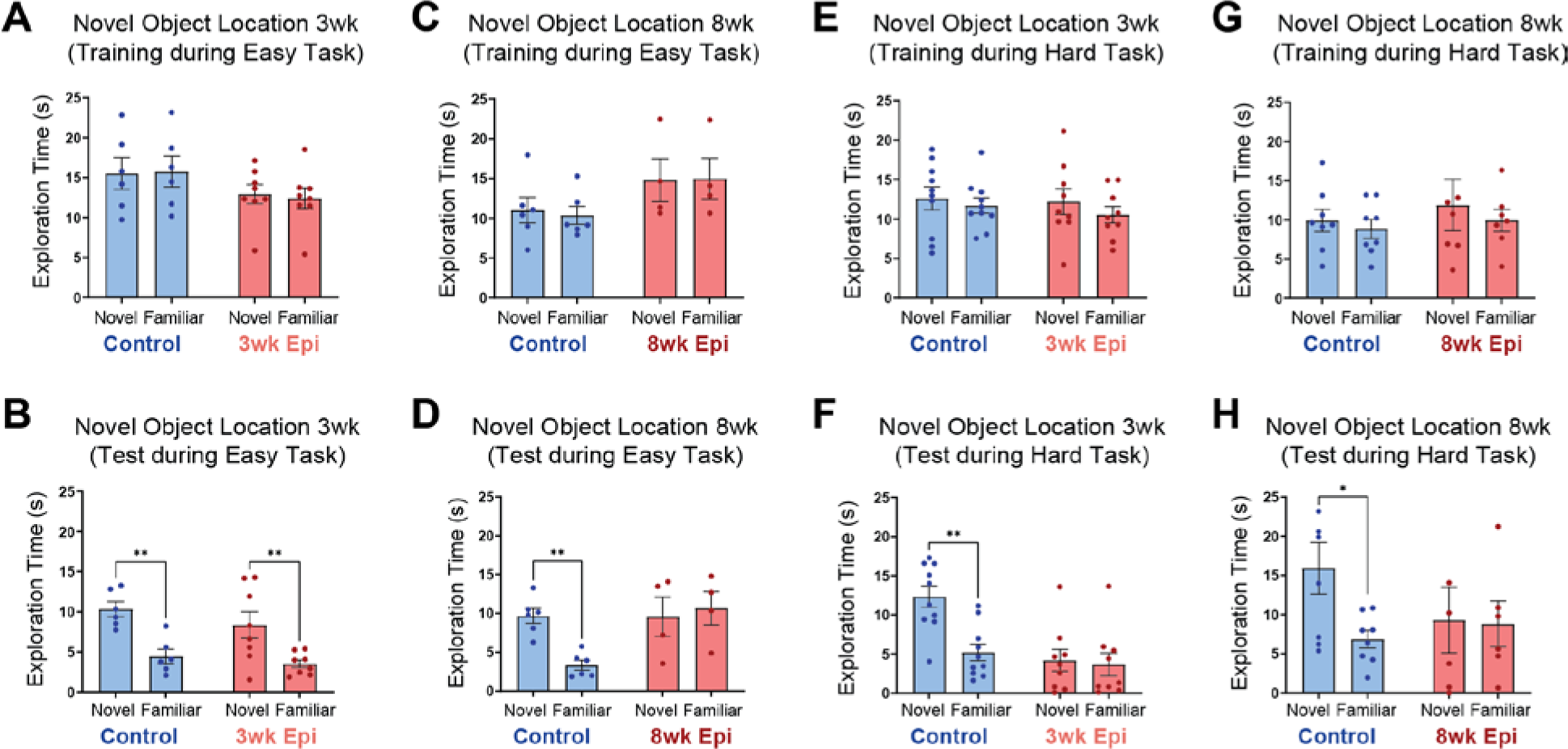
Novel object location (NOL) exploration times during training and testing. **A.** No differences in exploration time between the two objects in either Control or 3wk Epileptic groups during training on the Easy NOL task (N = 6, Control; N = 8, 3wk Epileptic; 2-way repeated measures ANOVA, *Fobject(1,12)* = 0.2, *p* > 0.05). **B.** Both Control and 3wk Epileptic groups spent more time with the moved object during testing on the Easy NOL task (N = 6, Control; N = 8, 3wk Epileptic; 2-way repeated measures ANOVA, *Fobject(1,12)* = 24.9, *p* < 0.001; Bonferroni corrected: Control: Novel v. Familiar *p* < 0.01; 3wk Epileptic: Novel v. Familiar *p* < 0.01). **C.** No differences in exploration time between the two objects in either Control or 8wk Epileptic groups during training on the Easy NOL task (N = 6, Control; N = 4, 8wk Epileptic; 2-way repeated measures ANOVA, *Fobject(1,8)* = 0.2, *p* > 0.05). **D.** During testing on the Easy NOL task, the Control group spent more time with the moved object while the 8wk Epileptic group spent equal time with both objects (N = 6, Control; N = 4, 8wk Epileptic; 2-way repeated measures ANOVA, *Fobject(1,8)* = 8, *p* < 0.05; Bonferroni corrected: Control: Novel v. Familiar *p* < 0.01; 8wk Epileptic: Novel v. Familiar *p* > 0.05). **E.** No differences in exploration time between the two objects in either Control or 3wk Epileptic group during training on the Hard NOL task (N = 10, Control; N = 9, 3wk Epileptic; 2-way repeated measure ANOVA, *Fobject(1,17)* = 2.5, *p* > 0.05). **F.** During testing on the Hard NOL task, the Control group spent more time with the moved object while the 3wk Epileptic group spent equal time with both objects (N = 10, Control; N = 9, 3wk Epileptic; 2-way repeated measure ANOVA, *Fobject(1,17)* = 8.5 *p* < 0.01; Bonferroni corrected: Control: Novel v. Familiar *p* < 0.01; 3wk Epileptic: Novel v. Familiar *p* > 0.05). **G.** No differences in exploration time between the two objects in either Control or 8wk Epileptic group during training on the Hard NOL task (N = 8, Control; N = 7, 8wk Epileptic; 2-way repeated measure ANOVA, *Fobject(1,13)* = 1.5, *p* > 0.05). **H.** During testing on the Hard NOL task, the Control group spent more time with the moved object while the 8wk Epileptic group spent equal time with both objects (N = 8, Control; N = 6, 8wk Epileptic; 2-way repeated measure ANOVA, *Fobject(1,12)* = 3.4, *p* = 0.09; Bonferroni corrected: Control: Novel v. Familiar *p* < 0.05; 8wk Epileptic: Novel v. Familiar *p* > 0.05). Error bars represent s.e.m. *p<0.05, **p<0.01, ***p<0.001

**Supplementary Figure 2:**
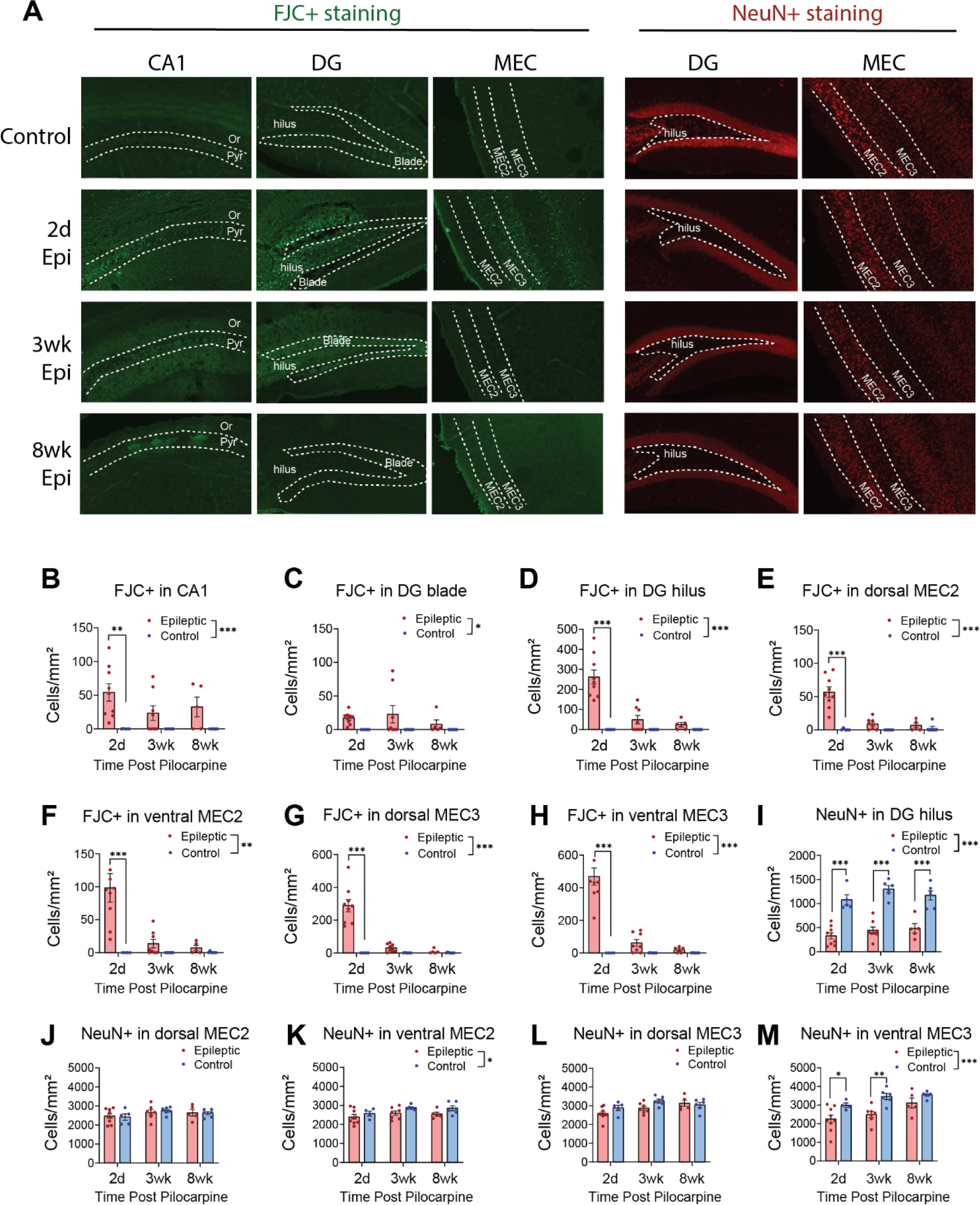
Neurodegenerative signaling and cell loss at 2 days, 3 weeks, and 8 weeks after Pilo-SE. **A.** Example immunohistochemistry staining for Fluro Jade-C (FJC, left) in CA1, DG (hilus and blade), MEC (MEC2 and MEC3), and NeuN (right) in DG (hilus) and MEC (MEC2 and MEC3) of Control and Epileptic mice. For Epileptic mice, tissue was collected at 2 days, 3 weeks, and 8 weeks after Pilo-SE. **B.** FJC staining was increased in CA1 in 2d Epileptic group, with group differences between Control and Epileptic groups (N = 5, 2d Control; N = 6, 3wk Control; N = 6, 8wk Control; N = 9, 2d Epileptic; N = 8, 3wk Epileptic; N = 5, 8wk Epileptic. 2-way ANOVA, *Fgroup(1,33)* = 18.2, *p* < 0.001; Bonferroni corrected: Control v. 2d Epileptic *p* < 0.01). **C.** FJC staining shows group level reduction in DG blade in epileptic groups (N = 5, 2d Control; N = 6, 3wk Control; N = 6, 8wk Control; N = 9, 2d Epileptic; N = 8, 3wk Epileptic; N = 5, 8wk Epileptic. 2-way ANOVA, *Fgroup(1,33)* = 7.2, *p* < 0.05). **D.** FJC staining was increased in DG hilus in the 2d Epileptic group, with a main effect between Control and Epileptic groups (N = 5, 2d control; N = 6, 3wk Control; N = 6, 8wk Control; N = 9, 2d Epileptic; N = 8, 3wk Epileptic; N = 5, 8wk Epileptic. 2-way ANOVA, *Fgroup(1,33)* = 32, *p* < 0.0001; Bonferroni corrected: Control v. 2d Epileptic *p* < 0.0001). **E.** FJC staining was increased in dorsal MEC2 in the 2d Epileptic group, with a main effect between Control and Epileptic groups (N = 5, 2d Control; N = 6, 3wk Control; N = 6, 8wk Control; N = 9, 2d Epileptic; N = 8, 3wk Epileptic; N = 5, 8wk Epileptic. 2-way ANOVA, *Fgroup(1,33)* = 29.3, *p* < 0.0001; Bonferroni corrected: Control v. 2d Epileptic *p* < 0.0001). **F.** FJC staining was increased in ventral MEC2 in the 2d Epileptic group, with a main effect between Control and Epileptic groups (N = 5, 2d Control; N = 6, 3wk Control; N = 6, 8wk Control; N = 9, 2d Epileptic; N = 8, 3wk Epileptic; N = 5, 8wk Epileptic. 2-way ANOVA, *Fgroup(1,33)* = 12, *p* < 0.01; Bonferroni corrected: Control v. 2d Epileptic *p* < 0.0001). **G.** FJC staining was increased in dorsal MEC3 in 2d Epileptic group, with a main effect between Control and Epileptic groups (N = 5, 2d Control; N = 6, 3wk Control; N = 6, 8wk Control; N = 9, 2d Epileptic; N = 8, 3wk Epileptic; N = 5, 8wk Epileptic. 2-way ANOVA, *Fgroup(1,33)* = 34.1, *p* < 0.0001; Bonferroni corrected: Control v. 2d Epileptic *p* < 0.0001). **H.** FJC staining was increased in ventral MEC3 in 2d Epileptic group, with a main effect between Control and Epileptic groups (N = 4, 2d Control; N = 6, 3wk Control; N = 6, 8wk Control; N = 9, 2d Epileptic; N = 8, 3wk Epileptic; N = 5, 8wk Epileptic. 2-way ANOVA, *Fgroup(1,32)* = 42.3, *p* < 0.0001; Bonferroni corrected: Control v. 2d Epileptic *p* < 0.0001). **I.** NeuN staining was decreased in DG hilus in all Epileptic groups, with a main effect between Control and Epileptic groups (N = 5, 2d Control; N = 6, 3wk Control; N = 6, 8wk Control; N = 9, 2d Epileptic; N = 8, 3wk Epileptic; N = 5, 8wk Epileptic. 2-way ANOVA, *Fgroup(1,33)* = 128.3, *p* < 0.0001; Bonferroni corrected: Control v. 2d Epileptic *p* < 0.0001; Control v. 3wk Epileptic *p* < 0.0001; Control v. 8wk Epileptic *p* < 0.0001). **J.** NeuN staining showed no difference in dorsal MEC2 between Epileptic and Control groups (N = 5, 2d Control; N = 6, 3wk Control; N = 6, 8wk Control; N = 9, 2d Epileptic; N = 7, 3wk Epileptic; N = 5, 8wk Epileptic. 2-way ANOVA, *Fgroup(1,32)* = 0, *p* > 0.05). **K.** NeuN staining was reduced in ventral MEC2 with a main effect between Control and Epileptic groups (N = 4, 2d Control; N = 6, 3wk Control; N = 6, 8wk Control; N = 9, 2d Epileptic; N = 7, 3wk Epileptic; N = 5, 8wk Epileptic. 2-way ANOVA, *Fgroup(1,31)* = 6.6, *p* < 0.05). **L.** NeuN staining showed no difference in dorsal MEC3 between Epileptic and Control groups (N = 5, 2d Control; N = 6, 3wk Control; N = 6, 8wk Control; N = 9, 2d Epileptic; N = 7, 3wk Epileptic; N = 5, 8wk Epileptic. 2-way ANOVA, *Fgroup(1,32)* = 3.7, *p* > 0.05). **M.** NeuN staining was decreased in ventral MEC3 in 2d and 3wk Epileptic groups, with a main effect between Control and Epileptic groups (N = 4, 2d Control; N = 6, 3wk Control; N = 6, 8wk Control; N = 9, 2d Epileptic; N = 8, 3wk Epileptic; N = 5, 8wk Epileptic. 2-way ANOVA, *Fgroup(1,31)* = 19, *p* < 0.001; Bonferroni corrected: Control v. 2d Epileptic *p* < 0.05; Control v. 3wk Epileptic *p* < 0.01). Error bars represent s.e.m. *p<0.05, **p<0.01, ***p<0.001.

**Supplementary Figure 3:**
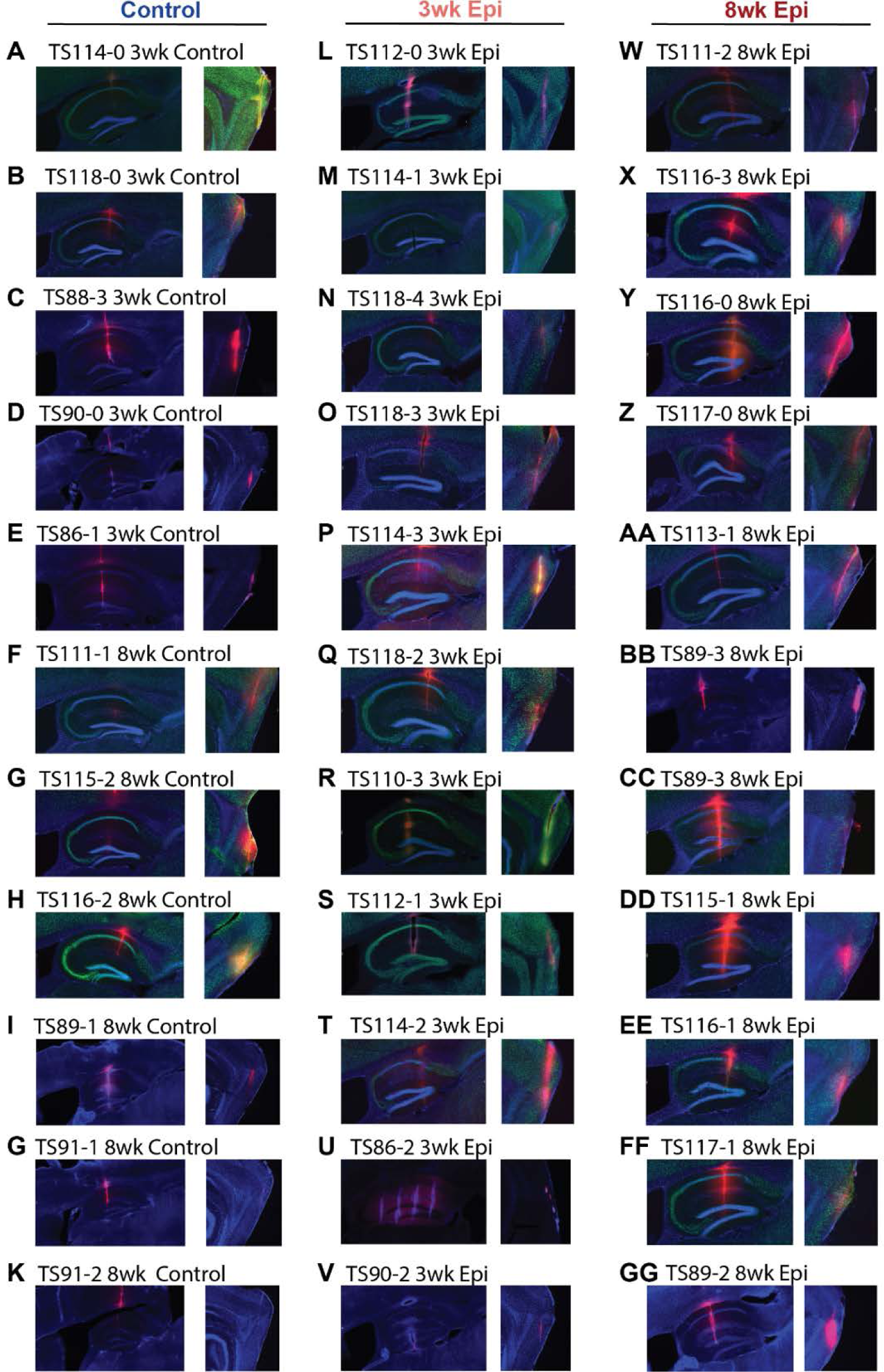
Representative probe tract for each animal. **A-K.** Probe tracts in all Control animals **L-V.** Probe tracts in all 3wk Epileptic (Epi) animals **W-GG.** Probe tracts in all 8wk Epileptic (Epi) animals Red: probe tract; Green: NeuN; Blue: DAPI

**Supplementary Figure 4:**
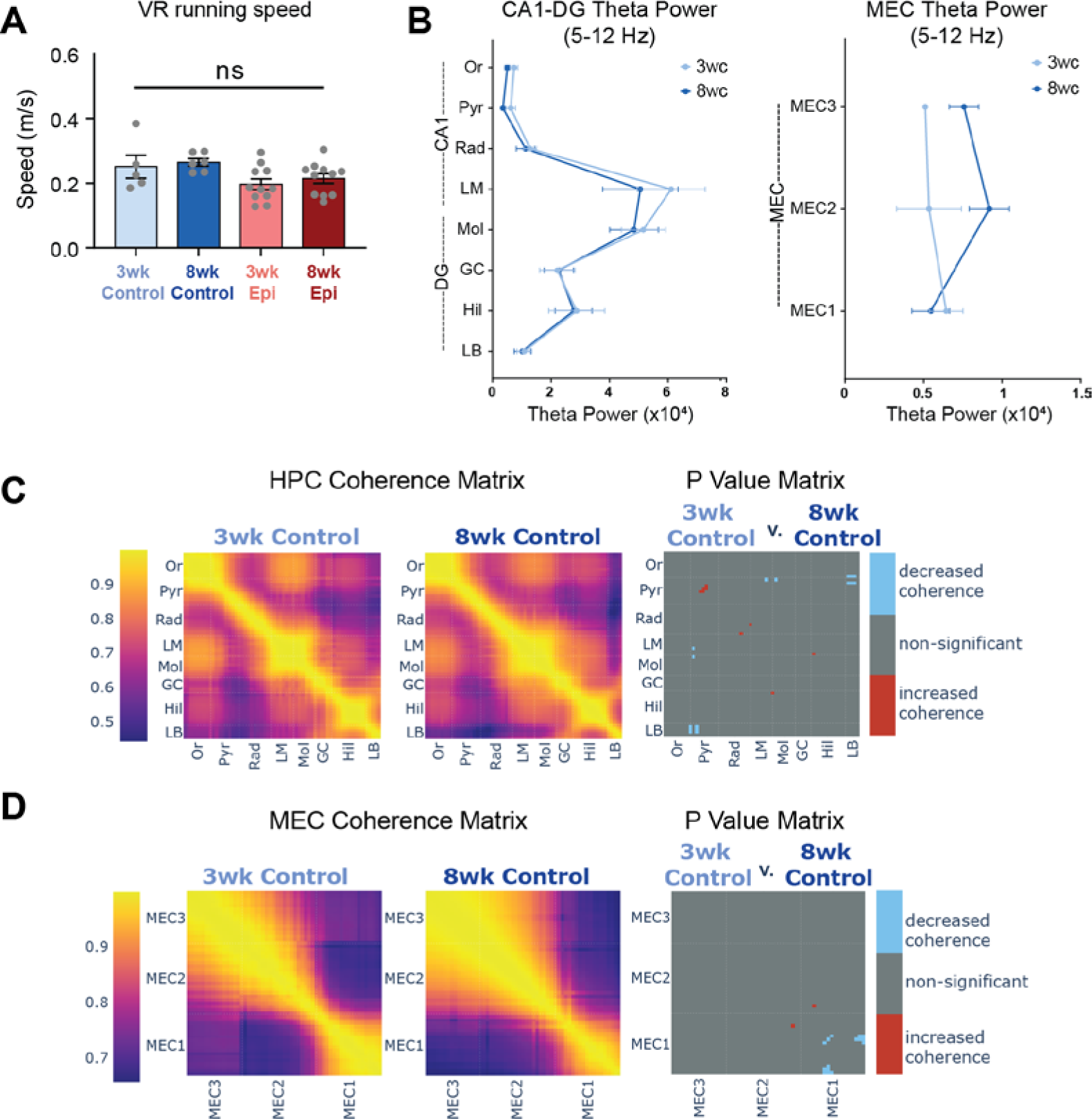
No differences in running speed across groups or theta power and coherence in Control animals. **A.** No differences in average running speed in VR during recordings (N = 5, 3wk Control; N = 6, 8wk Control; N = 11, 3wk Epileptic; N = 11, 8wk Epileptic. One-way ANOVA, *F(3,29)* = 2.6, *p* > 0.05). **B.** Theta power from each hippocampus layer (left) and MEC layer (right) in 3wk and 8wk Control animals. No differences were detected in any region (N = 5, 3wk Control; N = 6, 8wk Control; For HPC: Repeated-measures mixed-effects model, *F*Group x Region (7,61) = 0.2, *p* > 0.05; For MEC: Repeated-measures mixed-effects model, *F*Group x Region (2, 12) = 2.3, *p* > 0.05). **C.** Theta coherence between each channel pair along the probe in HPC in 3wk Control (left) and 8wk Control (middle) groups. P value matrix (right) shows the significant comparisons from each region pairs in HPC between groups. No clear patterns of significant differences were detected (N = 5, 3wk Control; N = 6, 8wk Control; welch t-test with alpha = 0.05; blue: *p* < 0.05, decrease coherence; red: *p* < 0.05, increase coherence). **D.** Theta coherence between each channel pair along the probe in MEC in 3wk Control (left) and 8wk Control (middle) groups. P value matrix (right) shows the significant comparisons from each region pairs in MEC between groups. No clear patterns of significant difference were detected (N = 5, 3wk Control; N = 6, 8wk Control; welch t-test with alpha = 0.05; blue: *p* < 0.05, decrease coherence; red: *p* < 0.05, increase coherence). Error bars represent s.e.m. *p<0.05, **p<0.01, ***p<0.001

**Supplementary Figure 5:**
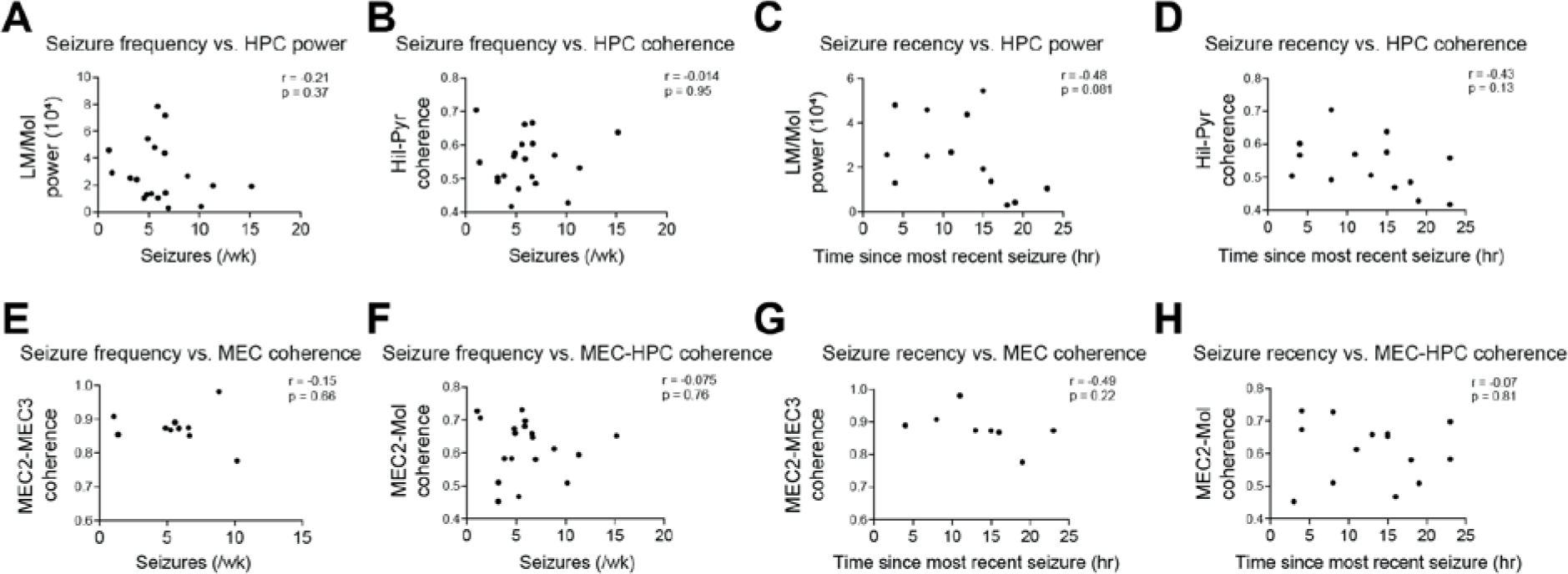
Seizure frequency and seizure recency is not correlated with theta power or coherence in Epileptic animals. **A.** No correlation between seizure frequency and HPC (LM and Mol layers) theta power (N = 20, Epileptic animals; Pearson r = −0.21, p > 0.05). **B.** No correlation between seizure frequency and HPC (Hil and Pyr layers) theta coherence (N = 20, Epileptic animals; Pearson r = −0.014, p > 0.05). **C.** No correlation between the time since the most recent seizure and theta power in HPC (LM and Mol layers) (N = 14, Epileptic animals; Pearson r = −0.48, p > 0.05). **D.** No correlation between the time since the most recent seizure and theta coherence between Hil and Pyr layers (N = 14, Epileptic animals; Pearson r = −0.43, p > 0.05). **E-H.** No correlation between seizure frequency and MEC (MEC2 and MEC3 layers) theta coherence (E: N = 11, Epileptic animals; Pearson r = −0.15, p > 0.05) or seizure frequency and MEC-HPC (MEC2 and Molecular layers) theta coherence (F: N = 19, Epileptic animals; Pearson r = −0.075, p > 0.05). No correlation between the time since the most recent seizure and theta coherence between layers MEC2 and MEC3 (G: N = 8, Epileptic animals; Pearson r = −0.49, p > 0.05) or theta coherence between MEC2 and HPC Molecular layer (H: N = 14, Epileptic animals; Pearson r = −0.07, p > 0.05).

**Supplementary Figure 6:**
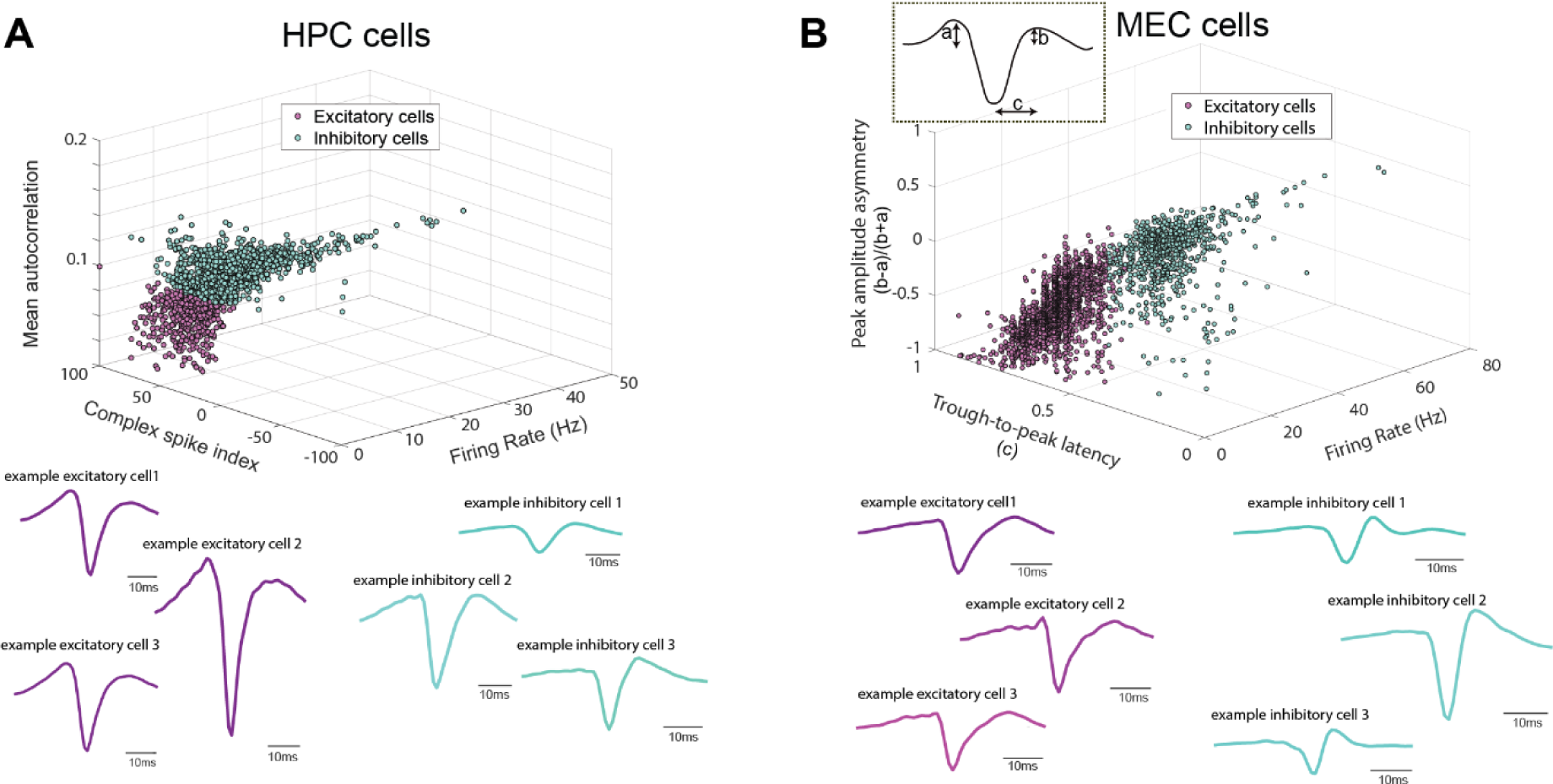
HPC and MEC putative excitatory and inhibitory cells. **A.** Top: Putative excitatory and inhibitory cells in HPC separated by mean autocorrelation, complex spike index, and firing rate. Bottom: 3 examples of excitatory HPC cells and 3 examples of inhibitory HPC cells. **B.** Top: Putative excitatory and inhibitory cells in MEC separated by trough-to-peak latency and peak amplitude asymmetry. Bottom: 3 examples of excitatory MEC cells and 3 examples of inhibitory MEC cells

**Supplementary Figure 7:**
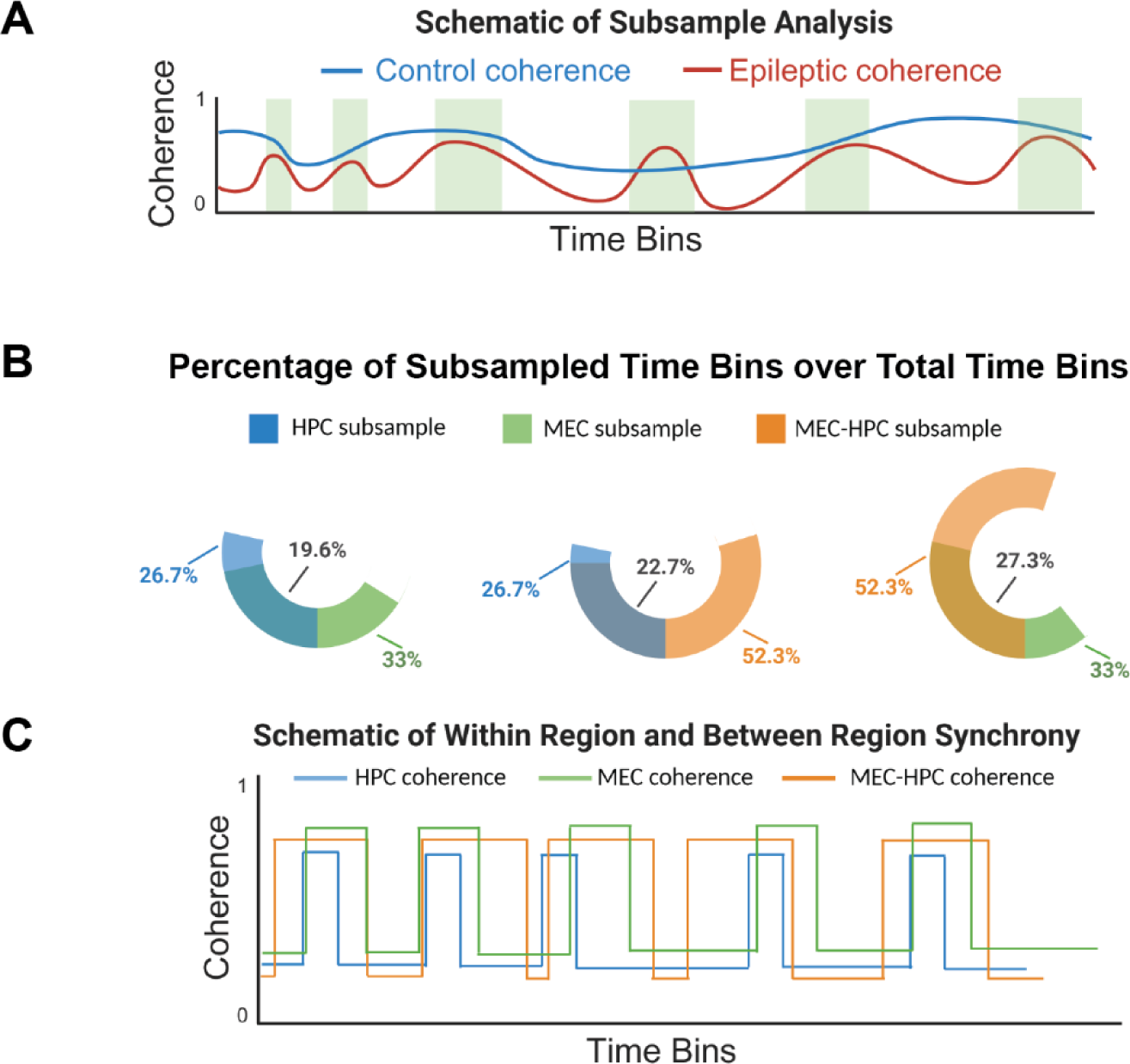
Schematic of subsample analysis and relationship between subsampled time bins. **A.** Schematic of subsample analysis. Blue: coherence level in Control animals; Red: coherence level in Epileptic animals; Green: time periods that Epileptic animals have the same level of coherence as in the Control group. Periods in green were selected for subsample analysis. **B.** Percent of time bins that were subsampled based on within-MEC, within-HPC, or MEC-HPC theta coherence, averaged across animals (HPC subsample: N = 11, 3wk Epileptic; N = 11, 8wk Epileptic; MEC subsample: N = 7, 3wk Epileptic; N = 5, 8wk Epileptic; MEC-HPC subsample: N = 10, 3wk Epileptic; N = 10, 8wk Epileptic). The percent of overlap in these time bins is shown in the darker shade. **C.** Schematic of the within-region and across-region coherence profile. During the time HPC coherence (blue) is high, MEC-HPC (orange) coherence is also high; during the time MEC-HPC (orange) coherence is high, not all HPC (blue) or MEC (green) coherence are high.

## Notes

### Competing Interest Statement

The authors have declared no competing interest.

